# CONVERGENCE, SAMPLING AND TOTAL ORDER ESTIMATOR EFFECTS ON PARAMETER ORTHOGONALITY IN GLOBAL SENSITIVITY ANALYSIS

**DOI:** 10.1101/2024.02.25.582013

**Authors:** Harry Saxton, Xu Xu, Torsten Schenkel, Richard H. Clayton, Ian Halliday

**Affiliations:** Materials & Engineering Research Institute Sheffield Hallam University, Sheffield, S1 1WB; Department of Computer Science, Insigneo Institute for in silico Medicine, University of Sheffield, Sheffield, S1 4DP; Department of Engineering and Mathematics Sheffield Hallam University, Sheffield, S1 1WB; Department of Infection, Immunity and Cardiovascular Disease, University of Sheffield, The Medical School, Sheffield, S10 2RX

**Keywords:** Sensitivity Analysis, Sobol’s Indices, Total Order Estimators, Modelling practices, Practical identifiability

## Abstract

Dynamical system models typically involve numerous input parameters whose “effects” and orthogonality need to be quantified through sensitivity analysis, to identify inputs contributing the greatest uncertainty. Whilst prior art has compared total-order estimators’ role in recovering “true” effects, assessing their ability to recover robust parameter orthogonality for use in identifiability metrics has not been investigated. In this paper, we perform: (i) an assessment using a different class of numerical models representing the cardiovascular system, (ii) a wider evaluation of sampling methodologies and their interactions with estimators, (iii) an investigation of the consequences of permuting estimators and sampling methodologies on input parameter orthogonality, (iv) a study of sample convergence through resampling, and (v) an assessment of whether positive outcomes are sustained when model input dimensionality increases. Our results indicate that Jansen or Janon estimators display efficient convergence with minimum uncertainty when coupled with Sobol and the lattice rule sampling methods, making them prime choices for calculating parameter orthogonality and influence. This study reveals that global sensitivity analysis is convergence driven. Unconverged indices are subject to error and therefore the true influence or orthogonality of the input parameters are not recovered. This investigation importantly clarifies the interactions of the estimator and the sampling methodology by reducing the associated ambiguities, defining novel practices for modelling in the life sciences.

**Research Highlights:**

- We conduct a heuristic investigation utilising 2 physiologically intuitive, highly nonlinear and stiff, lumped parameter models.
- The Janon and Jansen estimators emerge as optimal choices for calculating parameter orthogonality, as they are insensitive to sampling methodologies and measurement types.
- The Janon and Jansen estimators prove to have the most efficient convergence rates in calculating total order indices.
- The convergence rate of an estimator appears to be decisive in its ability to truthfully and uniformly recover true indices and orthogonality.
- Our methods provide putative best practice for practical identifiability investigations.

**Author Summary:** In order to gain a new insight into biological systems one often uses a mathematical model to predict possible responses from the system of interest. One vital step when using such models is knowledge of the uncertainty associated with a model response given a change in the inputs provided to the model. Utilising two non-linear and stiff cardiovascular models as test cases we investigate the effects of different choices made when quantifying the uncertainty in a mathematical model. Leveraging efficient solving of the mathematical model we are able to show that in order to truly quantify the effects of inputs on a set of outputs one must ensure converged estimates of the inputs influence. Without this, identifying inputs of a model become uncertain, or clinically, non patient specific. Our detailed study provides a workflow and advice for mathematical models of biological systems thus ensuring a true interpretation of the uncertainty associated with model inputs.

## 1 Introduction

Parameter identifiability addresses the question of whether, and to what degree it is possible to uniquely estimate input parameters (inputs) for a given dynamical system with a set of measured (or synthetic) outputs. This problem is typically decomposed into practical identifiability, which incorporates practical estimation issues associated with real data (such as noise and bias), and structural identifiability, which considers only model structure. The latter investigation is deemed ideal and effectively assumes that all data are known at every time point and are free of bias and noise [1]. Practical identifiability accounts for the role of noise and sampling frequency *inter alia* in hindering the ability uniquely to estimate inputs. These issues notwithstanding, the study of unique parameter estimation is very important to the complex models increasingly used in life sciences, which encompass pharmacology, epidemiology and cardiovascular applications [2, 3, 4]. Assuming one can identify inputs representative of the data, we arrive at *model personalisation*- a process of effectively calibrating a life science model using data available from an individual subject or patient. Within a clinical setting, this might involve calibrating a cardiovascular model to (patho)physiological metrics. Robust and reliable model personaisation is a key component for the development of digital twins for healthcare applications [5].

Unique model parameters are normally obtained by solving an inverse problem. One seeks the extrema of a cost function, typically the *L*_2_ norm of some weighted difference between measured, often very noisy and self-inconsistent target experimental data and a corresponding model prediction. A model cost function interacts with a hyper-surface in the model’s input parameter space and personalisation amounts to an attempt to locate the global minimum of the cost function. It is appropriate to emphasise, here, that input parameter space is multi-dimensional (with a dimension equalling the number of input parameters) and that the gradient of the cost function hyper-surface typically varies rapidly in some parameter directions (axes), but very slowly in others. Hence, a sufficient consideration of a model’s input parameter hyper-surface is central to understanding which parameters can be recovered uniquely.

Various methods exist to calculate model identifiability (see e.g [6]); our approach is based on the method of Li et al. [7], which calculates the *identifiability index* of the *i*th model input *I*_*i*_ as follows:

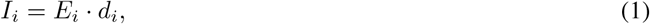

where *d*_*i*_ is the *i*th input’s orthogonality relative to a pre-selected, existing set of input parameters (which one is seeking to expand) and *E*_*i*_ is an *i*th input’s effect. Here, the index *I*_*i*_ measures the likelihood of a unique recovery of the *i*th model input. In our method, both effect and orthgonality of an input are calculated from the sensitivity matrix, generated with respect to model outputs [7, 8, 9, 10]. Clearly, the identifiability index depends on both the effect and the orthogonality which can be disclosed by sensitivity analysis. This prompts an investigation to find the most reliable and most robust method for calculation.

Sensitivity analysis studies how a change in a model’s specific output can be attributed to different sources of uncertainty in its (likely) many inputs [11]. Two types of sensitivity analysis exist: (i) Local sensitivity analysis (LSA), which examines the sensitivity of the model inputs at one specific model operating point in the input parameters’ space; (ii) Global sensitivity analysis (GSA), which determines sensitivities at multiple points throughout input space, before finding a statistic measure [12]. Variance-based indices are commonly recognised as the pre-eminent statistic of GSA, due to their model-independent nature, and their ability to account for interactions between model inputs and the ease of interpretation [13, 14]. For studies on practical identifiability of inputs, the metrics on the total order sensitivity matrix are calculated following Eq. (1), which evaluates the overall contribution of an input and its interaction with other inputs to a specific output whilst considering orthogonality. We defer further detailed discussion to Section 2.3.

Calculations of inputs’ effects and orthogonality are an important area of research for model personalisation. Here we ask three universal and interrelated questions:

i. What is the most reliable estimator for the underlying sensitivity indices to be computed on?
ii. What is the optimal sampling methodology, in relation to (i) above, for which one explores the complex input parameter space?
iii. How do the choices of estimator and sampling methodology impact the index convergence?

Previous work [15, 16, 17] mostly concentrates on efficient and accurate computation of the total order matrix and the evaluation of different estimators’ abilities to reveal the “true” effects of inputs, given their interactions. Recently, Puy et al. [18] reported their examination of several total order estimators - essentially a sensitivity analysis of a sensitivity analysis [19]. These authors varied the sampling method, between Monte Carlo and quasi-Monte Carlo, their analytic (note) test model, the dimensionality of input parameter hyperspace, the distribution of input parameters, and the number of model runs. The work provides a comprehensive and systematic assessment of the properties of different estimators.

We structure the paper as follows: Section 2 introduces the sampling methodologies and the total order estimators used for our investigation, as well as the nonlinear systems they are applied to; Section 3 presents our findings and section 4 declares and discusses best practice to offset interactions between sampling methodologies, estimators and model dimensionality when considering the orthogonality of inputs.

## 2 Methods

### 2.1 Backgound

Within the field of parameter identifiability, it is accepted [20] that there is interaction between the methodology of sampling from the model’s input parameter hyperspace and the estimators used to extract values of, e.g. Sobol indices [16]. It follows that other sensitivity derived metrics, such as the parameter orthogonality [7], are also affected by this same interaction. In this work, therefore, we extend the consideration of both sensitivity and orthogonality in tandem. We choose to undertake this assessment based upon a very important class of cardio-vascular models, which are notoriously stiff and non-linear, in contradistinction to the simple, traditional models usually used for such purposes [19, 21, 18], which possess analytic solutions. Because there is no analytical solution to the models considered in this work, the reference (or true) values of the sensitivity and orthogonality indices must therefore be determined using high levels of (re)sampling, based upon accepted processes of resampling and bootstrapping [21].

Our dynamical system has the form:

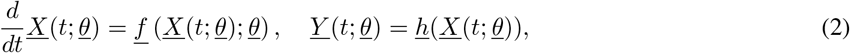

where <u>*Y*</u> (*t*; <u>*θ*</u>) represents the dynamic outputs described by the measurement functions, <u>*h*</u>, which depend on the state variables <u>*X*</u>(*t*; <u>*θ*</u>). The state variables are parameterised by the inputs <u>*θ*</u> but (for our application, at least) <u>*h*</u> does not directly depend upon <u>*θ. f*</u> represents a set of square integrable functions which, together with inputs, <u>*θ*</u>, determine <u>*X*</u>(*t*; <u>*θ*</u>), the solution of equations (2). Accordingly, <u>*f*</u> (<u>*X*</u>(*t*; <u>*θ*</u>); <u>*θ*</u>)), now viewed as a function of <u>*θ*</u>, is a suitable surrogate to determine the sensitivity of model inputs on outputs, through <u>*X*</u>(*t*; <u>*θ*</u>). For inputs which range over a bounded region, we consider a functional *I*[<u>*f*</u>](*t*), formed from the integral of <u>*f*</u> over the *n* dimensional hypercube *I*^*n*^, where *n* represents the dimensionality of the input parameter space:

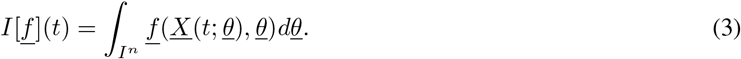

The effect of sampling inputs can be assessed with reference to this integral, by viewing the sampling process as an effective quadrature in which the sampled inputs define the abscissae. The quality of the sampling can then be measured by the quality of the quadrature. The important question of how its accuracy is determined in conjunction with the sampling of the hypercubic region of input parameter space is central to a robust and reliable sensitivity analysis.

### 2.2 Sampling Methodologies

In this section, we declare and briefly describe the input parameter space sampling methodologies that will be assessed in this work. We concern ourselves with two popular Monte Carlo (MC) sampling methodologies: Uniform (U) and Latin Hypercube (LH), and three Quasi-Monte Carlo (QMC) sampling methods: Golden Ratio (GR), Lattice Rule (LR) and Sobol Sequence (SS).

#### 2.2.1 Monte-Carlo Sampling Methods

##### Uniform sampling

The simplest sampling approach from literature is uniform sampling [22]. Input parameters <u>*θ*</u> are regarded as uniformly distributed random variables, within the hypercube *I*^*n*^ such that:

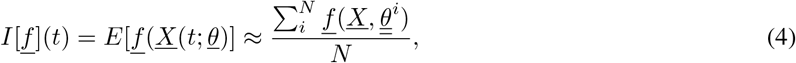

where 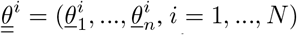 is a sequence of independent random points in *I*^*n*^ of length *N*. This is deemed a crude approximation with poor convergence rates [23].

##### Latin hypercube sampling

The efficiency of MC methods is determined by the properties of the random samples. A priority for researchers is to develop strategies which ensure points are placed more uniformly, within *I*^*n*^. One response is to use LH [24], which is a very common methodology in life sciences [25, 26, 27, 28]. Its main objective is to reduce the variance associated with evaluating Eq. (3). One decomposes the space of inputs into *N* -dimensional squares, to ensure the space is sampled as uniformly as possible. Let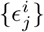, for *j* = 1, …, *n*, be independent random permutations of samples *i* = 1, …, *N*, each uniformly distributed over all *N* ! possible permutations. One then sets

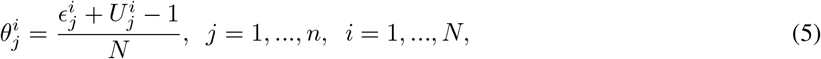

where 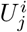 are independently randomly sampled points on the [0,1] interval. It can be seen intuitively how only one sample point of the input parameters falls between 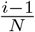 and 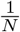 for each dimension *j* = 1, …, *n*. Here, we use the ‘standard’ version of LH, but see [29] for other variations.

#### 2.2.2 Quasi Monte-Carlo Sampling Methods

An improvement on the Monte-Carlo sampling methodologies is the low discrepancy sampling (LDS) methods, coupled with the QMC algorithm, as shown in [30, 31]. Discrepancy is a measure of the deviation of sampled points from the uniform distribution [32]. Consider a number of points *N*_*R*_ from a sequence {*θ*_*i*_}, for *i* = 1, .., *N*, in an *n*-dimensional rectangle *R* centred upon an origin 0, whose sides are parallel to the coordinate axis, which is a subset of *I*^*n*^ : *R* ⊂ *I*^*n*^, where *R* is attached with a measure. A sequence has low discrepancy if the proportion of points in the sequence falling into an arbitrary set *R* is close to the measure of *R*. LDS satisfies the upper bound condition [33]:

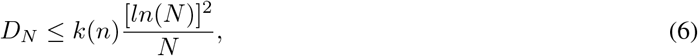

where *D*_*N*_ is the sample discrepancy and *k*(*n*) is a particular constant depending on the sequence and size of input paraeter space. LDS is designed to place sample points as uniformly as possible mathematically, within a hypercube, instead of the statistical approach adopted in LH. The QMC approximation of the integral in Eq. (3) has identical form to Eq. (4).

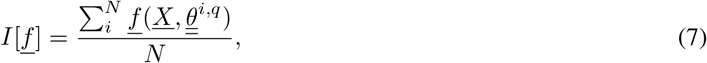

except in this framework, 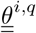 is a matrix which has been generated from an LDS and the points are distributed uniformly in the hypercube *I*_*n*_. As a consequence, the sample points generated on *I*^*n*^ have a deterministic nature.

##### Golden ratio sampling

Golden ratio sampling is an LDS sampler in which sample points are based on the fractional part of successive integer multiples of the golden ratio. First introduced by Schretter and Kobbelt [34], using a simple incremental permutation of a generated golden ratio sequence, they demonstrated equal coverage of a two-dimensional space. The generating sequence is expressed as:

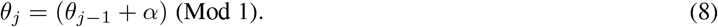

The constant *α* that gives the lowest possible discrepancy is 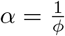, where *ϕ* is the golden ratio [35]. *j* is the counter for each parameter sampled. In this work and specifically in Section 3, we will test GR sampling for systems with input parameter dimensions much higher than two.

##### Rank-1 lattice rule sampling

Another LDS is the rank-1 lattice rule, where an *n*-dimensional rank-1 lattice Π is a set of points that contains no limit points and satisfies [36]:

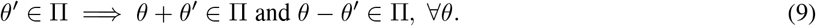

A general lattice is constructed by a generating matrix *G* ∈ ℝ^*n×n*^:

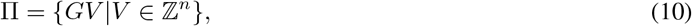

where *V* is any integer unimodular integer vector. A generator matrix is not unique to a lattice Π, i.e., Π can be obtained from different generator matrices. A rank-1 lattice is a special case of the general lattice, which has a simple operation for point set construction, instead of directly using Eq. (10). A rank-1 lattice point set can be constructed as

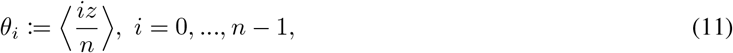

where *z* ∈ ℤ is the generating vector and the inner product denotes the operation of taking the fractional part of the input number elementwise. Compared with the general lattice rule, the construction form of the rank-1 lattice already ensures the constructed points are inside the unit cube without the need for any further checks.

##### Sobol sequence sampling

Our final sampling methodology is the well-known Sobol LDS [37]. The Sobol sequence is widely considered as the optimal sequence for exploration of an input parameter space [27, 38, 39, 40]. The construction of the Sobol sequence uses linear recurrence relations over the finite field 𝔽. Let the binary expansion of the non-negative integer *R* be given by 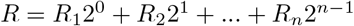, where *n* is the dimension of the input parameter hypercube. Then the *j*th element of the *n*th dimension of the Sobol sequence, 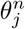can be generated by:

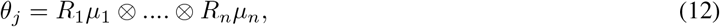

where 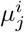 is a binary fraction called the *i*th direction number in the *j*th dimension. These direction numbers are generated by the following *q*-term recurrence relation:

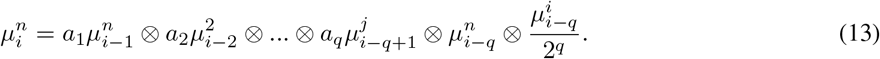

We have *i > q*, and *a*_*i*_ comes from the coefficients of a degree-*q* primitive polynomial over 𝔽. Put simply, the Sobol LDS aims to achieve three requirements: (1) Best uniformity as *n* → ∞; (2) Good distribution even with small parameter sizes; (3) A very fast computational algorithm.

### 2.3 Sensitivity Analysis and Orthogonality

Given a model of the form in Eq. (2) with *Y* (a continuous or discrete output), a variance based first order or total order effect can be calculated for a generic input factor *θ*_*i*_. 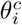 denotes the complementary set, i.e., all other model inputs excluding *θ*_*i*_. The first order sensitivity index can be written as:

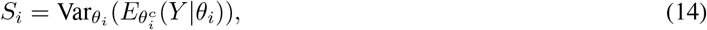

where *E* is the expectation operator. The inner expectation operator functions such that the mean of *Y* is taken over all possible values of 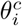 while keeping *θ*_*i*_ fixed. The outer variance is taken over all possible values of *θ*_*i*_. Then utilising the known identity [41]:

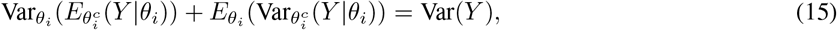

where 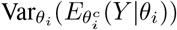 measures the first order (additive) effects of *θ*_*i*_ on the model outputs. Another popular variance measure (and the concentration of this work) is total order estimators, first introduced by Homma [42]:

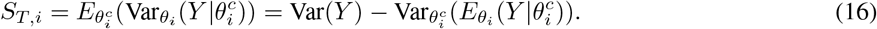

Here *S*_*T,i*_ measures the total effect, i.e., first and higher order effects (multiplicative interactions) of input parameter *θ*_*i*_. One can consider this by recognising that 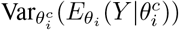 is the first order effect of 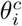, so 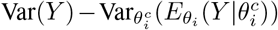 must give the contribution of all terms in the variance decomposition which do include the input *θ*_*i*_.

The equations can be derived through a Hoeffding-Sobol decomposition, and utilising the fact that each term is assumed to be square integrable. The detailed derivation can be found in [43, 44].

Sobol indices converge slowly in general and estimators are used to accelerate the process [11, 13]. Here, we benchmark five sampling methodologies against four commonly chosen total order estimators: Homma and Saltelli [42], Sobol [45], Jansen [46] and Janon et al. [47]. While this list is far from exhaustive, it represents a selection of total order estimators which have been practically used within the field and are not costly to execute computationally.

For the Homma & Saltelli, Sobol and Janson estimators, their mean and variance take the following form:

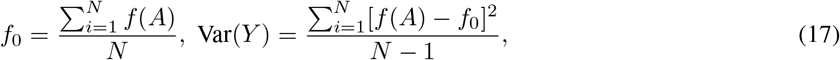

and for the Janon estimator:

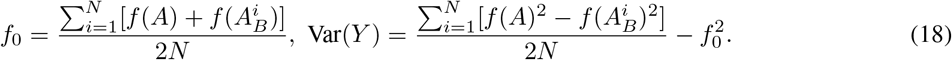

In order to utilise the above formulae, we propose two independent sampling matrices are generated - *A* and *B*, with elements *a*_*ij*_ and *b*_*ij*_, for *i* = 1, …, *N, j* = 1, …, *n* (where *N* is the number of samples and *n* is the total number of input parameters). We can now introduce a matrix 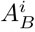 or 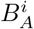 where all the rows are from *A* or *B*, except the *i*-th row which is extracted from *B* or *A*. These matrices are then used to compute the sensitivity indices which will be discussed below. While one can use either combination of the *A* and *B* matrices, within this work we use the couple *A*, 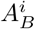 due to its proven efficiency in calculating sensitivity indices [16].

As will be defined in the next section, we will calculate total order indices for both continuous and discrete measurements. For continuous measurements, calculating the total order index produces waveform data which demonstrate the sensitivity of each input parameter over the cardiac cycle. In order to quantify the effects continuous measurements have on the calculation of total order, we must average this sensitivity waveform. Rather than averaging across a time range (which process regions of low variance equally to those of high variance), we seek to expose differential sensitivities by examining variance-weighted averages:

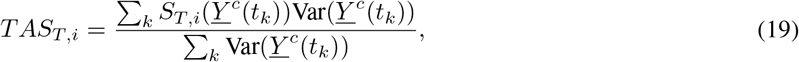

where *TAS*_*T,i*_ is the time averaged total order effect of an input parameter *i* and *Y* ^*c*^(*t*_*k*_) represents the approximated continuous measurement at time step *k*.

We use the measure *d*_*ij*_ to measure orthogonality between input parameters *θ*_*i*_ and *θ*_*j*_:

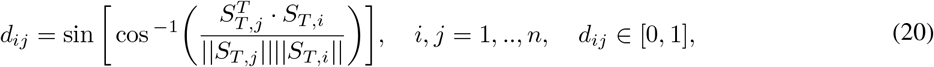

where 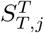 represents the transposed total order sensitivity matrix. Calculating this returns a *n × n* matrix where each element *d*_*ij*_ ∈ [0, 1]. *d*_*ij*_ = 0 represents total dependence between input parameters and *d*_*ij*_ = 1 represents complete independence. We then are able to rank input parameters by calculating the mean of input parameters’ orthogonality and ranking them such that the input parameter with the highest orthogonality is in position 1 and the input parameter with the lowest orthogonality is in position *n*, where *n* is the input space dimension.

### 2.4 Models and Data

We examine two models, which are representative of the heart and systemic circulation, of varying dimensionality in order to assess their potential effects on total order estimators. The first model is a three-compartment system-level, differential algebraic equations (DAE) based, electrical analogue cardiovascular (CV) model after Bjordalsbakke et. al. [48] which we will refer to as the **1-chamber model** (see Figure 1A). Our second model is a five-compartment model, introduced by Shi et al. [49], which we shall denote the **2-chamber model** (see Figure 1B). All model simulation code is avilable at https://github.com/H-Sax/Orthgonality-SA. The latter model is of the same form as Bjordalsbakke’s, however, the input parameter space dimensionality has increased from nine to twenty, due to the addition of an atrium and other compartments.

**Figure 1.**
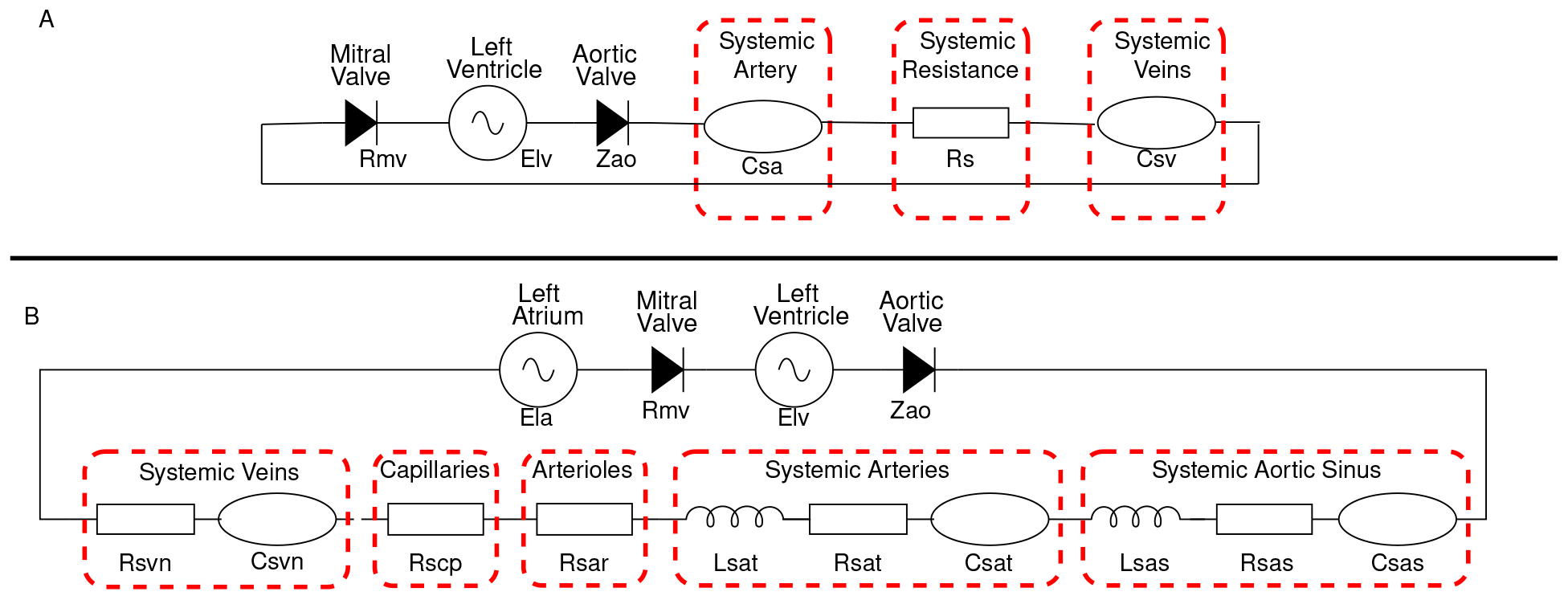
The two electrical analougue models utilised in this work. **(A)** is a nine parameter representation of the systemic circulation originally presented by Bjordalsbakke et al. [48]. **(B)** is a twenty parameter representation of the systemic circulation originally presented by Shi et al. [49].

Each compartment state is specified by its dynamic pressure *P* (*t*) (mmHg), an inlet flow *Q*(*t*) (mL/s) and a volume *V* (*t*) (mL):

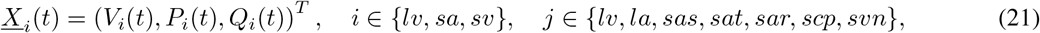

where for the 1 chamber model *lv, sa, sv* mean the left ventricle, systemic artery and systemic veins respectively. For the 2 chamber model *lv, la, sas, sat, sar, scp, svn* represent the left ventricle, left atrium, systemic aortic sinus, systemic artery, systemic arterioles, systemic capillary and systemic vein. Formally, *t* is the continuous variable time. For a full model description along with model parameters, see appendix A.

As we focus on the computation of total order indices, along with varying model dimensionality, we will also vary the type of data provided to the model between continuous and discrete. For the simple nine parameter model defined in Figure 1A, we utilise:

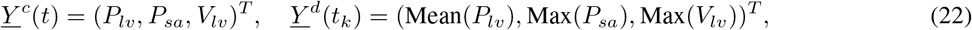

where *t*_*k*_ represents the discrete time point where the measurement is taken, <u>*Y*</u>^*c*^ and <u>*Y*</u>^*d*^ represent the continuous and discrete measurement vector. For the twenty dimensional model in Figure 1B we utilise similar measurements:

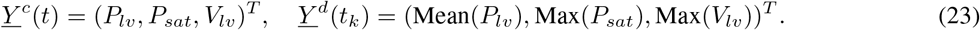

We follow the advice of Saltelli et al. [16] that the first and total order indices require *k* computations:

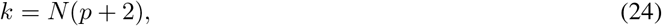

where *p* is the dimensionality of the input parameter space, *N* is the number of samples taken from the input parameter space and *k* is the total number of model evaluations needed in order to compute the indices. It is well accepted within the literature, that utilising the low discrepancy Sobol sequence combined with the Jansen Estimator is considered best practice for calculation of total order indices [16, 18].

Given the high non-linearity and stiffness associated with these lumped parameter models, we first perform an assessment of convergence and uncertainty in the calculation of the total order indices. We consider the indices converged once there is no more change in the rank of the input parameters and the error associated with the indices is less than 5%. Once the indices have converged with the Jansen estimator and Sobol sampling methodology, we fix this sample size, *N*, for all other estimators and sampling methodologies. For the 9-parameter, 1-chamber model shown in Figure 1A, we investigate the convergence by varying *N* ∈ [2000, 40000]. For the 20-parameter, 2-chamber model shown in Figure 1B, we vary *N* ∈ [10000, 30000]. The uncertainty associated with the indices is calculated using re-sampling with replacement, where we set the number of bootstraps to *B* = 1000, as found in the literature [21, 50]. Once convergence has been achieved for the discrete measurements, we use this sample size *N* for the continuous outputs to ensure the time averaged indices derived are not subject to excessive uncertainty.

All computations are performed using Julia [51] and reproducible code is available at https://github.com/H-Sax/Orthgonality-SA. Figure 2 details the workflow needed to generate the sensitivity indices in Julia. Step 1 involves defining the ODEs of the system of interest. Model A is defined employing the package DifferentialEquations.jl [52] and Model B is implemented using our acausal modelling library CirculatorySystemModels.jl [53]. Step 2 generates the sample points needed to perform GSA. QuasiMonteCarlo.jl is a Julia package which provides the needed sampling algorithms. Step 3 uses GlobalSensitivity.jl [54] which is a Julia implementation of the Sobol indices with varying estimators and bootstrapping methodology. Then to analyse the sensitivity indices, we utilise self written functions.

**Figure 2.**
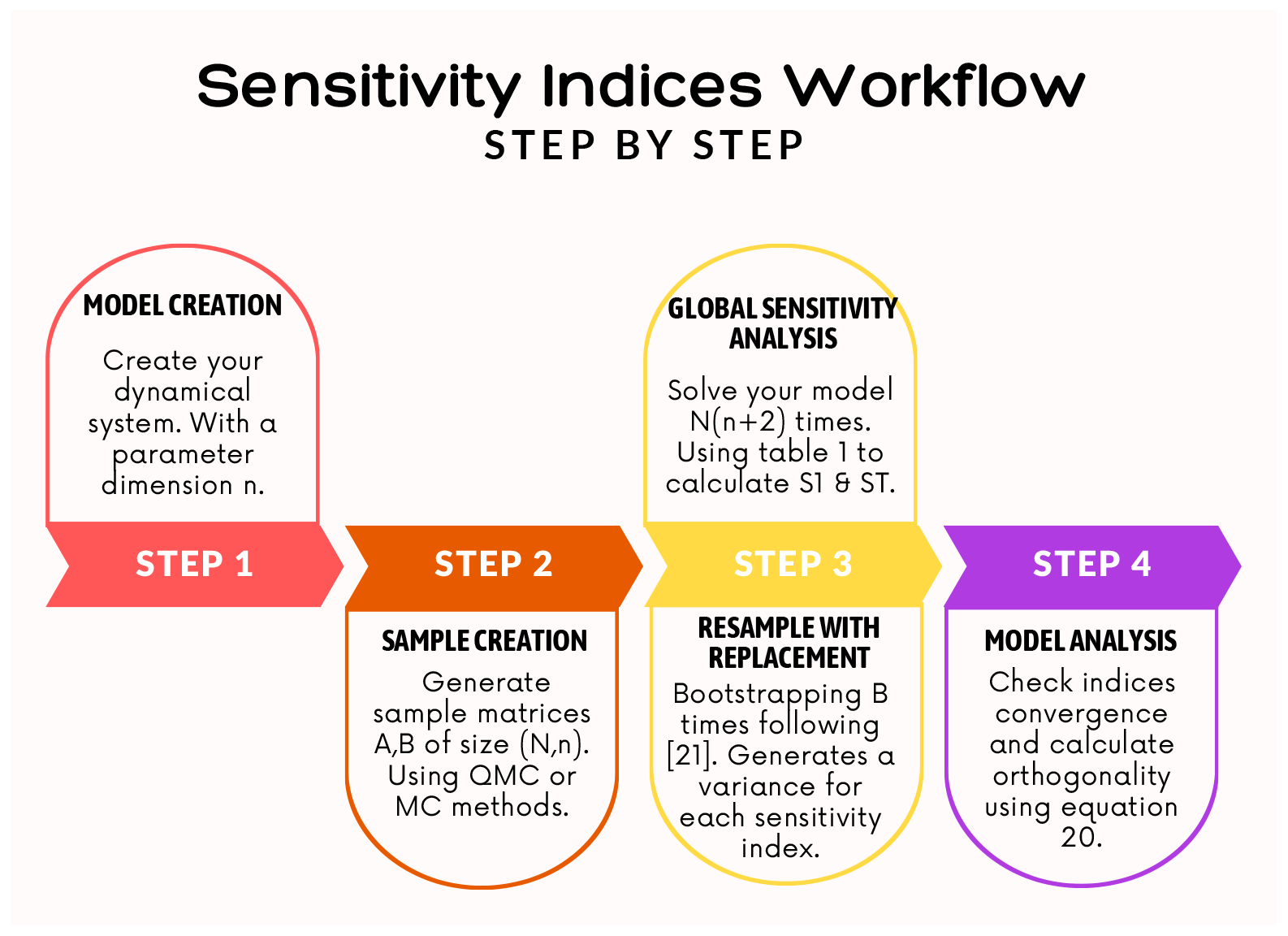
Workflow to compute sensitivity indices within Julia. Full code utilising these steps can be found at https://github.com/H-Sax/Orthgonality-SA.

Specifically, simulations were solved using Vern7 algorithm [55], with relative and absolute tolerances set to 10^−8^. We saved the model solution at 200 points between cycles 15 and 16, a steady solution being reached after 5 cycles and used Makie.jl to visualise results [56].

## 3 Results

In this section, we present: (i) the convergence of total order indices with respect to both discrete and continuous output measurements; (ii) our investigation outcomes of varying the four estimators with the five sampling methodologies defined in Section 2.2. First, the convergence results will be illustrated in Section 3.1. Then in the next two subsections, total order Sobol indices and orthogonality of input parameters for the 1-chamber, 9-parameter model (Figure 1A) and the 2-chamber, 20-parameter model (Figure 1B) are shown. Between each subsection, we examine what effect the different choices of estimator and sampling methodology has on the orthogonality of input parameters. Within each of the subsections, we make the distinction between the effects of continuous and discrete measurements. We present results for input parameters which are deemed to have high clinical significance (i.e., bio-markers), for example, low arterial compliance *C*_*sa*_ may indicate a stiffening of the vessel. Each subsection displays convergence of a single parameter for brevity.^2^. Physiologically realistic time series solutions generated from the model can be found in appendix A, figure 12.

### 3.1 Convergence and Uncertainty

Figure 3 shows the convergence and the uncertainty of the minimal ventricular elastance *E*_*min*_ for the 1-chamber and 2-chamber models. We calculate the Sobol indices using the Jansen estimator and Sobol sampling, which are considered to be best practice [18, 46]. Henceforth, they are regarded as our benchmark and the sample size returned from this initial investigation is used for evaluations on all other estimators and sampling methods. Increasing the sample size and re-sampling with replacement allow us to evaluate the sample size at which the Sobol indices have converged with minimum uncertainty. This is displayed as a band around the index of interest and represents a 95% confidence interval of the index estimate.

**Figure 3.**
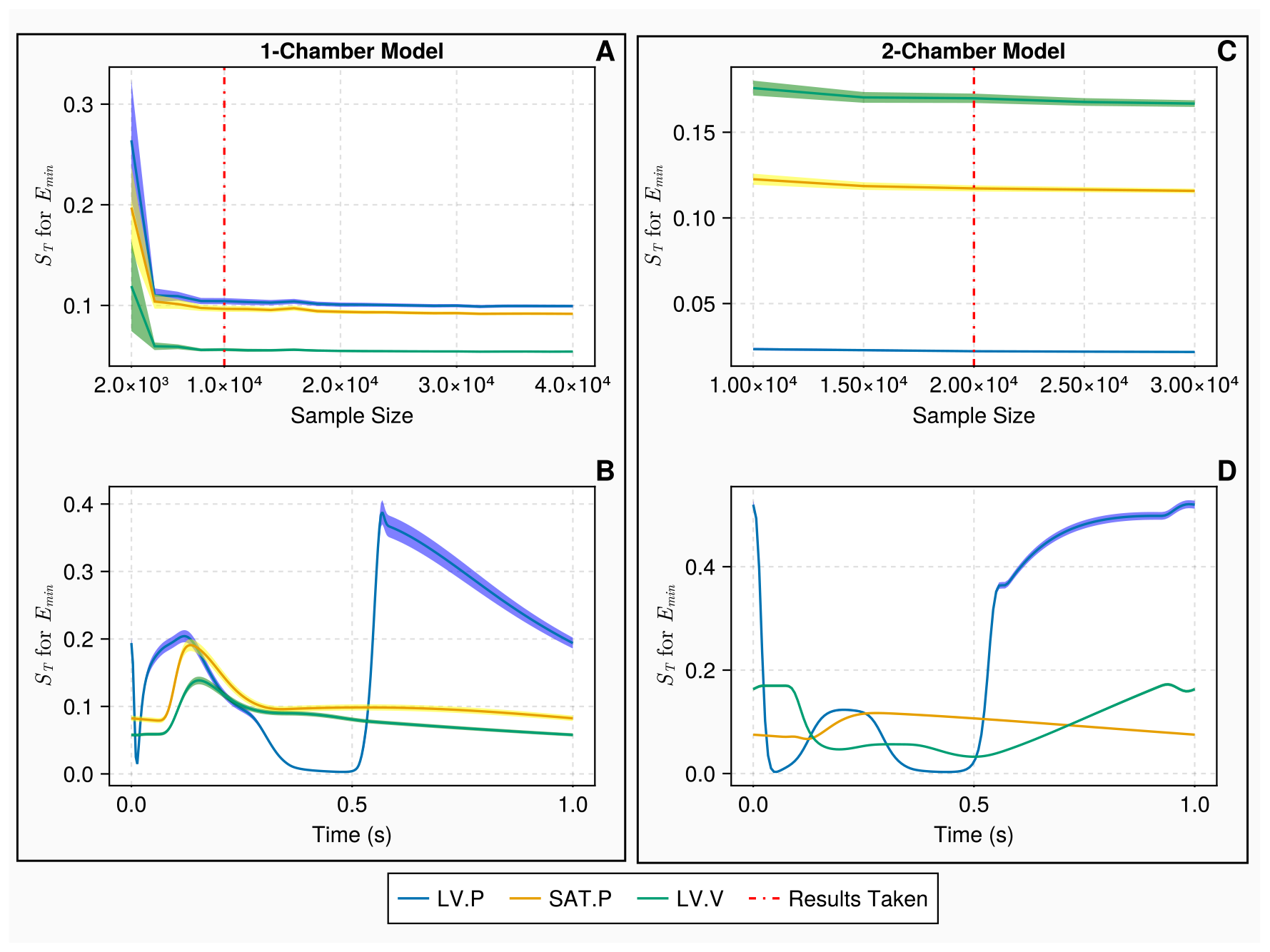
Convergence and uncertainty of indices associated with the minimum ventricular elastance *E*_*min*_. Figure **A** displays the convergence and uncertainty of the Sobol indices *S*_*T*_ calculated on discrete measurements for the 1-chamber model against increasing sample size. Here, the vertical line signifies the chosen sample size for the 1-chamber model at *N* = 10, 000. Figure **B** presents the continuous Sobol indices with uncertainty bounds, calculated at a sample size *N* = 10, 000, on continuous measurements over a single cardiac cycle, for the 1-chamber model. Figure **C** displays the convergence and uncertainty of *S*_*T*_ calculated on discrete measurements for the 2-chamber model against increasing sample size. Again, the vertical line signifies the chosen sample size for this model, at *N* = 20, 000. Figure **D** shows the continuous Sobol indices with uncertainty bounds for *N* = 20, 000, on continuous measurements over a single cardiac cycle, for the 2-chamber model. The measurements shown in blue, yellow and green denote the left ventricular pressure, the systemic arterial pressure and the left ventricular volume, respectively. In the discrete settings (i.e., A and C), the measurements are the mean left ventricular pressure, the maximum systemic arterial pressure and the maximum left ventricular volume.

For the 1-chamber model, 10, 000 samples (110, 000 model evaluations) ensured convergence when computed against the discrete measurements defined in Eq. (22). Figure 3A shows that evaluating the Sobol indices at a higher sample size would provide minimal improvement and at 10, 000 samples the indices are subject to negligible error. Figure 3B shows that the continuous indices have minimal error during the cardiac cycle so when we compute the time averaged indices no excessive error will be present. Using the 10, 000 sample size, we computed the continuous Sobol indices against the measurements defined in Eq. (22). For the 2-chamber model, 20, 000 samples (660, 000 model evaluations) were adequate as seen in Figure 3C. The continuous measurements of the 2-chamber model, seen in Figure 3D, show that the indices were not subject to error for 20, 000 samples. Figure 3C appears to indicate that fewer samples may be adequate for the 2-chamber model. However, the adopted sample size ensured all input parameters displayed a consistent rank with less than 5% error.

### 3.2 1-chamber Model

The uncertainty associated with the computation of Sobol total order indices on the 1-chamber model, with *N* = 10, 000 samples, is presented in Figure 4. Only the results for arterial compliance *C*_*sa*_ are displayed in this figure. The 95% confidence intervals for the Homma and Sobol estimators are considerably wider than that of the Jansen and Janon estimators. The Homma estimator consistently produced estimates of the sensitivity indices which are different to that of the other available estimators. The Jansen and Janon estimators are identical in their computations of the sensitivity indices and confidence intervals, invariant of the sampling methodology used. When the Homma and Sobol estimators are used, the latin hypercube and uniform sampling methods produce larger confidence intervals compared to the quasi-monte carlo sampling methods.

**Figure 4.**
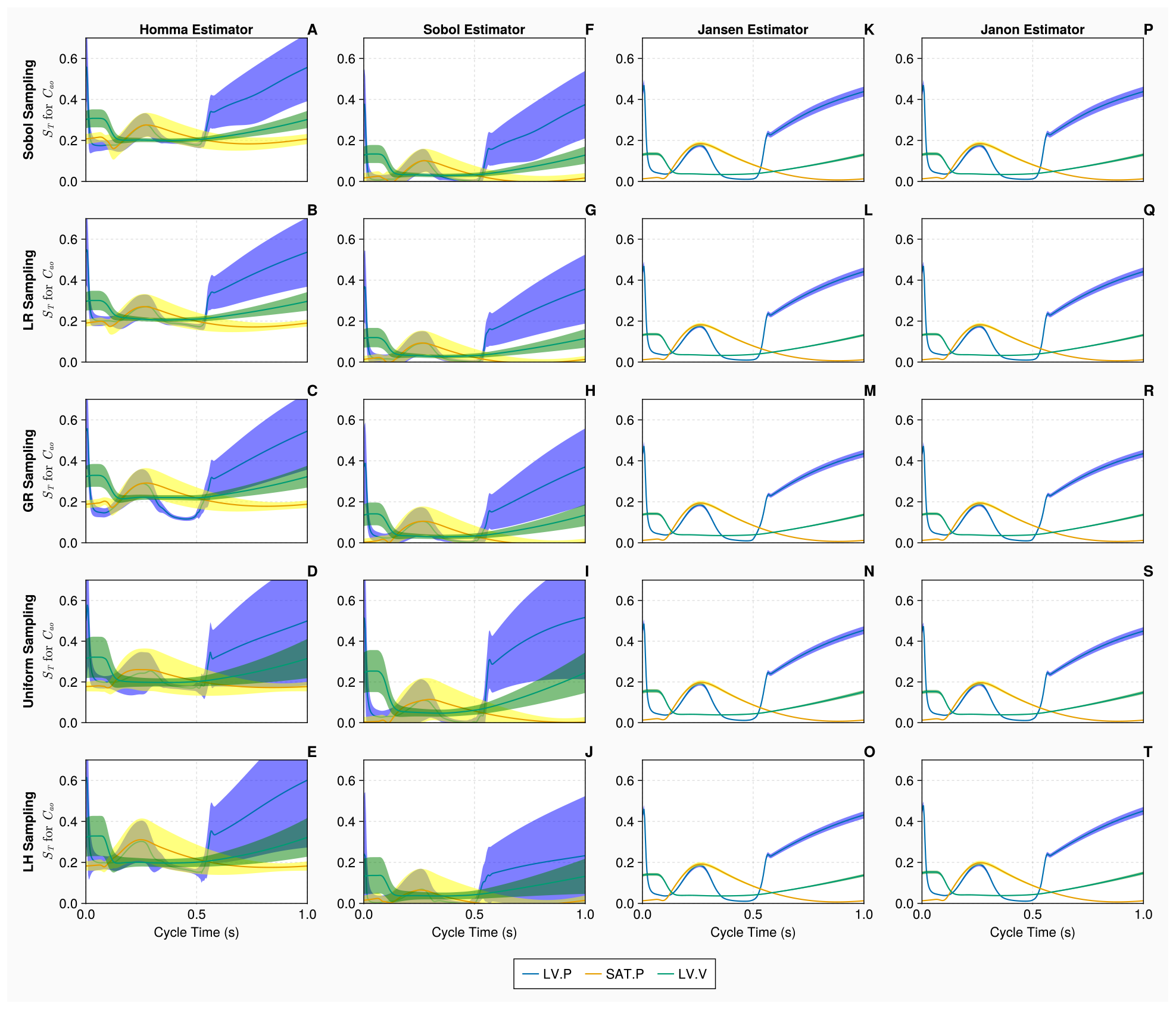
Total order Sobol indices *S*_*T*_ of the arterial compliance *C*_*sa*_ for the 1-chamber model with continuous measurements. Panels A - T show *S*_*T*_ of *C*_*sa*_, for 3 continuous measurements - left ventricular pressure, systemic arterial pressure and the left ventricular volume (represented in blue, yellow and green curves, respectively), over a single cardiac cycle with differing estimators and sampling methodologies. Measurements are evaluated with *N* = 10, 000 samples, using *B* = 1000 bootstrapped samples to evaluate the uncertainty of the estimate. The bands represent 95% confidence intervals associated with specific indices displayed as solid curves.

When calculating total order indices on the 1-chamber model with continuous measurements (the histograms for the orthogonality distributions of input parameters are presented in Figure 5), we notice that the orthogonality spreads for the Jansen and Janon estimators are identical for the golden ratio and Latin hypercube sampling methodologies. The Jansen estimator coupled with lattice rule sampling also shares this orthogonality distribution. The Janon estimator, with Lattice Rule and Sobol sampling, and the Jansen estimator with Sobol sampling, exhibit minor variations from the previous orthogonality distributions generated by Janon and Jansen estimators, however, are identical between themselves. The orthogonality distributions returned from the Homma and Sobol estimators exhibit large variations for each sampling methodology, although the Sobol estimator with the Sobol sampling returns an orthogonality distribution similar to that seen by the Jansen and Janon estimators. These results are mirrored in Table 2 where the input parameters are ranked based on their orthogonality scores in the parameter space. We see the orthogonality results obtained for the Jansen and Janon estimators are invariant to sampling methodologies. In contrast, the rankings for Sobol and Homma estimators vary amongst different sampling methodologies. The Sobol estimator when coupled with Sobol sampling, returns a parameter ranking almost identical to that of the Jansen and Janon estimators, a result consistent with the one observed in Figure 5.

**Table 1:** Formulae to compute *S*_*T,i*_, where *f*_0_ and Var represent the mean and variance of the outputs respectively, as defined in Eqs. (17) and (18).

| Authors | Estimator $S_{T,n}$ |
| --- | --- |
| Homma & Saltelli [42] | $\text{Var}(Y) - \frac{\sum_{i=1}^N f(A)f(A_B^i)}{N} + f_0^2$ |
| Sobol [45] | $\frac{\sum_{i=1}^N f(A)[f(A) - f(A_B^i)]}{N}$ |
| Jansen [46] | $\frac{\sum_{i=1}^N [f(A) - f(A_B^i)]^2}{2N}$ |
| Janon et al.[47] | $\text{Var}(Y) - \frac{\sum_{i=1}^N f(A)f(A_B^i)}{N} - f_0^2$ |

**Table 2:** Input parameter ranking for the 1-chamber model with continuous measurements. Here, input parameters are ranked based on the averaged orthogonality score returned from the calculated total order sensitivity matrix. In addition, the ranking is stratified by both sampling and estimator types.

| | | $\tau_{es}$ | $\tau_{ep}$ | $R_{mv}$ | $Z_{ao}$ | $R_s$ | $C_{sa}$ | $C_{sv}$ | $E_{max}$ | $E_{min}$ |
| --- | --- | --- | --- | --- | --- | --- | --- | --- | --- | --- |
| Homma | SS | 5 | 4 | 9 | 3 | 2 | 8 | 7 | 6 | 1 |
|  | LR | 5 | 4 | 9 | 3 | 2 | 8 | 6 | 7 | 1 |
|  | GR | 5 | 4 | 9 | 3 | 1 | 8 | 5 | 6 | 2 |
|  | U | 7 | 4 | 9 | 3 | 1 | 8 | 5 | 6 | 2 |
|  | LH | 6 | 4 | 8 | 1 | 3 | 9 | 5 | 7 | 2 |
| Sobol | SS | 4 | 3 | 8 | 5 | 1 | 7 | 9 | 6 | 2 |
|  | LR | 4 | 3 | 9 | 6 | 1 | 8 | 7 | 5 | 2 |
|  | GR | 4 | 3 | 8 | 7 | 1 | 6 | 9 | 5 | 2 |
|  | U | 5 | 3 | 8 | 4 | 1 | 9 | 7 | 6 | 2 |
|  | LH | 5 | 4 | 6 | 3 | 1 | 8 | 9 | 7 | 2 |
| Jansen | SS | 4 | 3 | 7 | 5 | 1 | 8 | 9 | 6 | 2 |
|  | LR | 4 | 3 | 7 | 5 | 1 | 8 | 9 | 6 | 2 |
|  | GR | 4 | 3 | 7 | 5 | 1 | 8 | 9 | 6 | 2 |
|  | U | 4 | 3 | 7 | 5 | 1 | 8 | 9 | 6 | 2 |
|  | LH | 4 | 3 | 7 | 5 | 1 | 8 | 9 | 6 | 2 |
| Janon | SS | 4 | 3 | 7 | 5 | 1 | 8 | 9 | 6 | 2 |
|  | LR | 4 | 3 | 7 | 5 | 1 | 8 | 9 | 6 | 2 |
|  | GR | 4 | 3 | 7 | 5 | 1 | 8 | 9 | 6 | 2 |
|  | U | 4 | 3 | 7 | 5 | 1 | 8 | 9 | 6 | 2 |
|  | LH | 4 | 3 | 7 | 5 | 1 | 8 | 9 | 6 | 2 |

**Figure 5.**
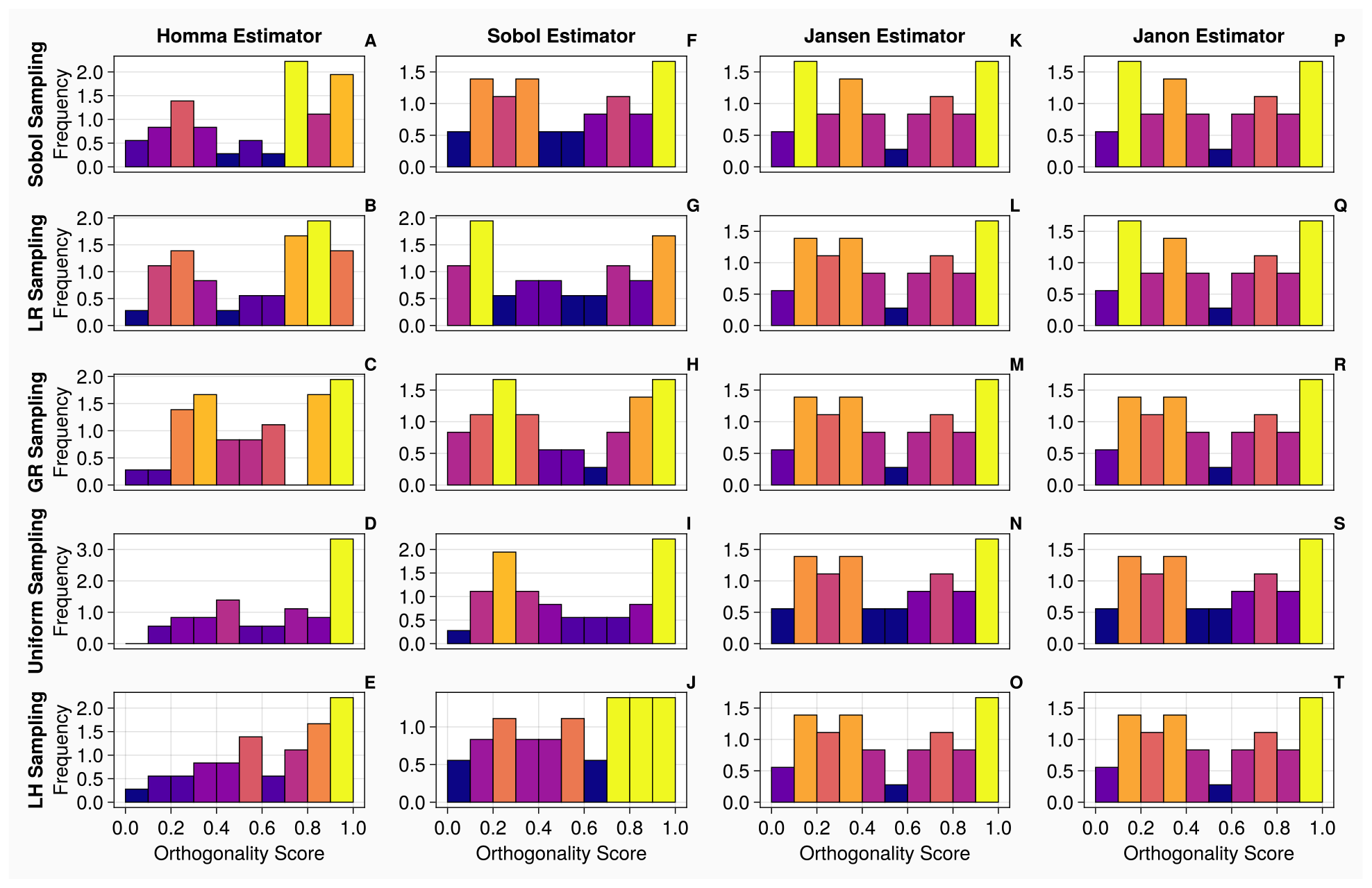
Orthogonality distributions of input parameters for the 1-chamber model with continuous measurements. Histograms A-T show the distribution of orthogonality returned from examinations of the sensitivity vectors, calculated from continuous measurements. Here, an orthogonality score of 1 represents total independence of input parameters, whereas 0 represents total dependence. Each individual diagram denotes a specific combination of sampling methodology and estimator type. The frequency of each histogram is normalised such that it is comparable between plots, i.e., the larger the frequency of a bin, the larger the number of orthogonality scores calculated from the original sensitivity vectors.

In Table 3, stratification by estimator type and examination of the range of an input parameter across all sampling methodologies reveal, as inferred from Table 2, that the Jansen and Janon estimators exhibit no variation for the whole input parameter set, given any sampling methodology. This indicates that the Jansen and Janon estimators are the optimal choices for this model. The Homma and Sobol estimators exhibit variations of 1.33 and 1.67 upon the input parameter set, respectively. These variations mean that using Homma and Sobol estimators will return differing orthogonality rankings when different sampling methodologies are used. When stratifying by sampling types, Table 4 reveals that Sobol and Lattice Rule samplings exhibit the smallest mean variations of the input set across all estimator types. It is important to note that these variations are a consequence of the Sobol and Homma estimators which both exhibited different orthogonality rankings for input parameters. These results indicate that given a less than optimal estimator, the Sobol or Lattice rule sampling methodology may produce a ranking which can be considered closer to the ground “truth”. Interestingly, we notice that the commonly used Latin Hypercube sampling methodology in life sciences exhibits the largest variation of an input set of parameters.

**Table 3:**
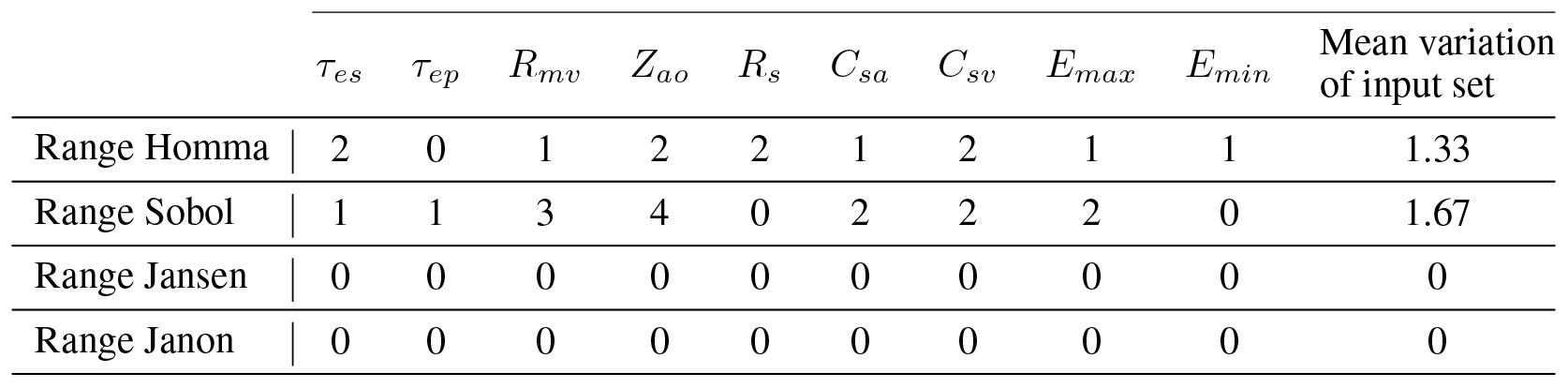
The ranges of input parameters across 5 sampling types for a specific estimator for the 1-chamber model with continuous measurements.

**Table 4:** The ranges of input parameters across 4 estimator types for a specific sampling method for the 1-chamber model with continuous measurements.

| | $\tau_{es}$ | $\tau_{ep}$ | $R_{mv}$ | $Z_{ao}$ | $R_s$ | $C_{sa}$ | $C_{sv}$ | $E_{min}$ | $E_{max}$ | Mean variation of input set |
| --- | --- | --- | --- | --- | --- | --- | --- | --- | --- | --- |
| Range SS | 1 | 1 | 2 | 2 | 1 | 1 | 2 | 0 | 1 | 1.22 |
| Range LR | 1 | 1 | 2 | 2 | 1 | 1 | 2 | 0 | 1 | 1.22 |
| Range GR | 1 | 1 | 2 | 4 | 0 | 2 | 3 | 2 | 0 | 1.67 |
| Range U | 3 | 1 | 1 | 2 | 0 | 2 | 4 | 0 | 0 | 1.44 |
| Range LH | 2 | 1 | 2 | 4 | 2 | 2 | 4 | 1 | 0 | 2.00 |

Figure 6 displays the convergence and uncertainty associated with the computation of the total order indices of the mitral value resistance *R*_*mv*_ for the 1-chamber model against the discrete measurements. In all cases, as the sample sizes are increased, the accuracy of the estimations and uncertainty associated with the indices improve. The Jansen and Janon estimators provide the most efficient convergences and smallest errors when calculating the indices. A sample size of *N* = 10, 000 is taken for the discrete measurements, because the columns for the Jansen and Janon estimators (Panels K - T) illustrate that any additional samples would return minimal improvements in terms of accurate calculation of the indices. The Homma and Sobol estimators (Panels A - J) display considerably larger errors than that of the Jansen and Janon estimators. When the upper limit sample size of *k* = 40, 000 is reached, the Homma and Sobol estimators appear to have converged with reduced errors when combined with the Sobol, Lattice Rule and Golden sampling methods, although the errors are still much larger than those exhibited by the Jansen and Janon estimators. The uniform and Latin hypercube sampling methods present the largest errors when combined with the Sobol estimator.

**Figure 6.**
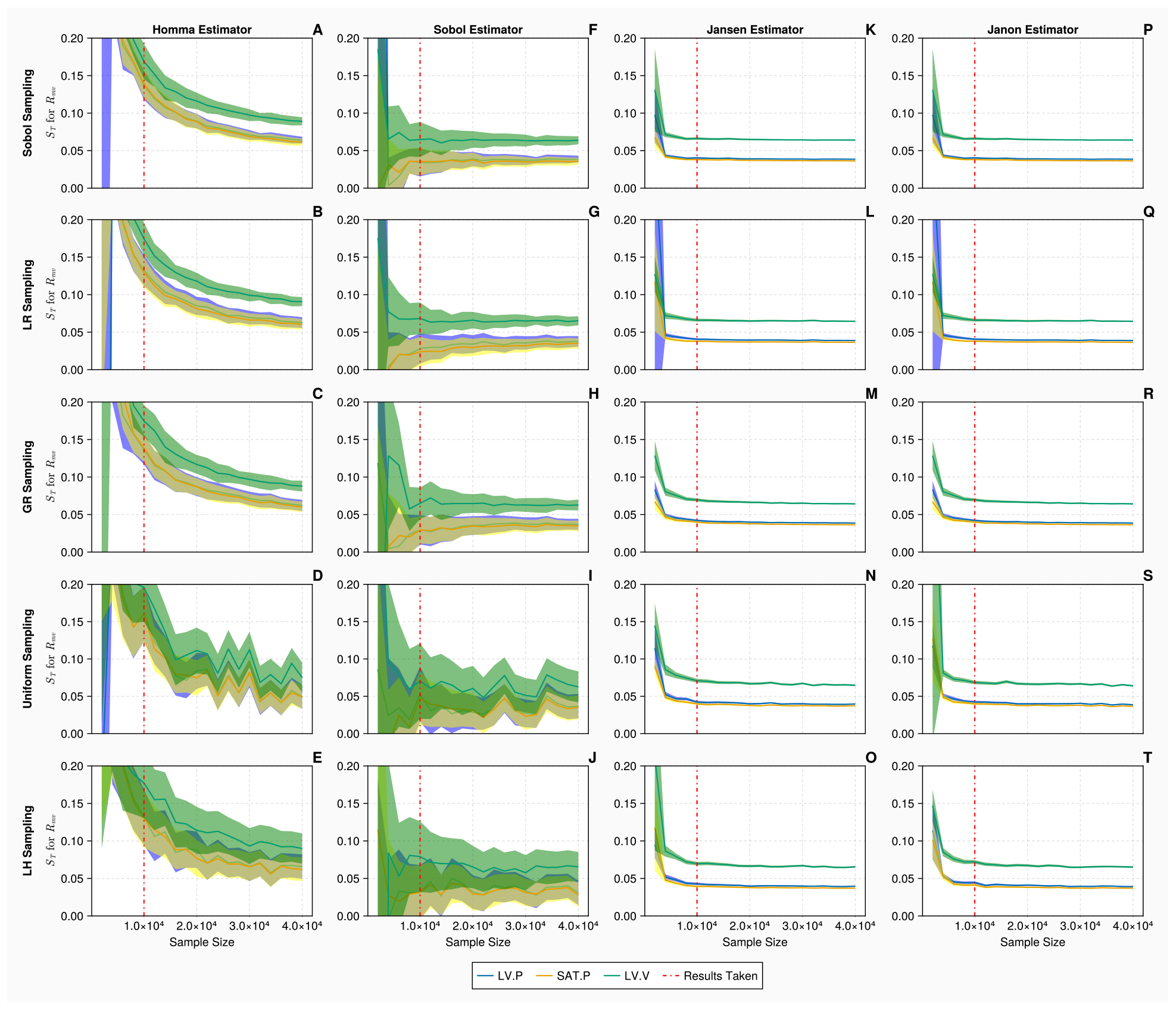
Total order Sobol indices *S*_*T*_ of the mitral valve resistance *R*_*mv*_ for the 1-chamber model with discrete measurements. Panels A - T show *S*_*T*_ of *R*_*mv*_, for 3 discrete measurements: mean left ventricular pressure, maximum systemic arterial pressure and maximum left ventricular volume (represented in blue, yellow and green, respectively), evaluated at increasing sample sizes (*N*∈ [2000, 40000] using *B* = 1000 bootstrapped samples), with differing estimators and sampling methodologies. The bands represent 95% confidence intervals associated with specific indices displayed as solid curves. The red solid vertical lines represent the point (*N* = 10, 000) at which the sample size is taken.

When calculating total order indices on the 1-chamber model with discrete measurements, the histograms presented in Figure 7 show that the orthogonality spreads for the Jansen and Janon estimators for all sampling methodologies, except the Janon estimator and the Latin Hypercube sampling pairing, are identical. We notice, the orthogonality distributions returned from the Homma estimator exhibit large variations for each sampling methodology, as seen with continuous measurements shown in Figure 5. The Sobol estimator column (Panels F - J) in Figure 7 displays orthogonality spreads which are somewhat similar to that of Jansen and Janon, however are still quite variant amongst sampling methodologies. These results are reflected in Table 5 where the Jansen and Janon estimators are shown to invariant to sampling methodologies, whilst the rankings of parameter orthogonality for the Sobol and Homma estimators vary amongst different sampling methodologies.

**Table 5:** Input parameter ranking for the 1-chamber model with discrete measurements. Again, input parameters are ranked based on the averaged orthogonality score returned from the calculated total order sensitivity matrix. The ranking is also stratified by both sampling and estimator types.

| | | $\tau_{es}$ | $\tau_{ep}$ | $R_{mv}$ | $Z_{ao}$ | $R_s$ | $C_{sa}$ | $C_{sv}$ | $E_{max}$ | $E_{min}$ |
| --- | --- | --- | --- | --- | --- | --- | --- | --- | --- | --- |
| Homma | SS | 3 | 1 | 4 | 2 | 5 | 9 | 8 | 6 | 7 |
|  | LR | 3 | 1 | 4 | 2 | 6 | 7 | 9 | 8 | 5 |
|  | GR | 2 | 4 | 6 | 1 | 7 | 5 | 8 | 9 | 3 |
|  | U | 5 | 2 | 8 | 6 | 7 | 3 | 9 | 4 | 1 |
|  | LH | 3 | 4 | 5 | 6 | 1 | 8 | 2 | 7 | 9 |
| Sobol | SS | 3 | 2 | 4 | 1 | 5 | 8 | 7 | 6 | 9 |
|  | LR | 6 | 2 | 4 | 1 | 3 | 9 | 7 | 5 | 8 |
|  | GR | 3 | 2 | 5 | 1 | 6 | 7 | 8 | 4 | 9 |
|  | U | 1 | 3 | 5 | 2 | 4 | 8 | 7 | 6 | 9 |
|  | LH | 7 | 1 | 4 | 2 | 3 | 6 | 8 | 5 | 9 |
| Jansen | SS | 5 | 2 | 3 | 1 | 4 | 9 | 7 | 6 | 8 |
|  | LR | 5 | 2 | 3 | 1 | 4 | 9 | 7 | 6 | 8 |
|  | GR | 5 | 2 | 3 | 1 | 4 | 9 | 7 | 6 | 8 |
|  | U | 5 | 2 | 3 | 1 | 4 | 9 | 7 | 6 | 8 |
|  | LH | 5 | 2 | 3 | 1 | 4 | 9 | 7 | 6 | 8 |
| Janon | SS | 5 | 2 | 3 | 1 | 4 | 9 | 7 | 6 | 8 |
|  | LR | 5 | 2 | 3 | 1 | 4 | 9 | 7 | 6 | 8 |
|  | GR | 5 | 2 | 3 | 1 | 4 | 9 | 7 | 6 | 8 |
|  | U | 5 | 2 | 3 | 1 | 4 | 9 | 7 | 6 | 8 |
|  | LH | 5 | 2 | 3 | 1 | 4 | 9 | 7 | 6 | 8 |

**Figure 7.**
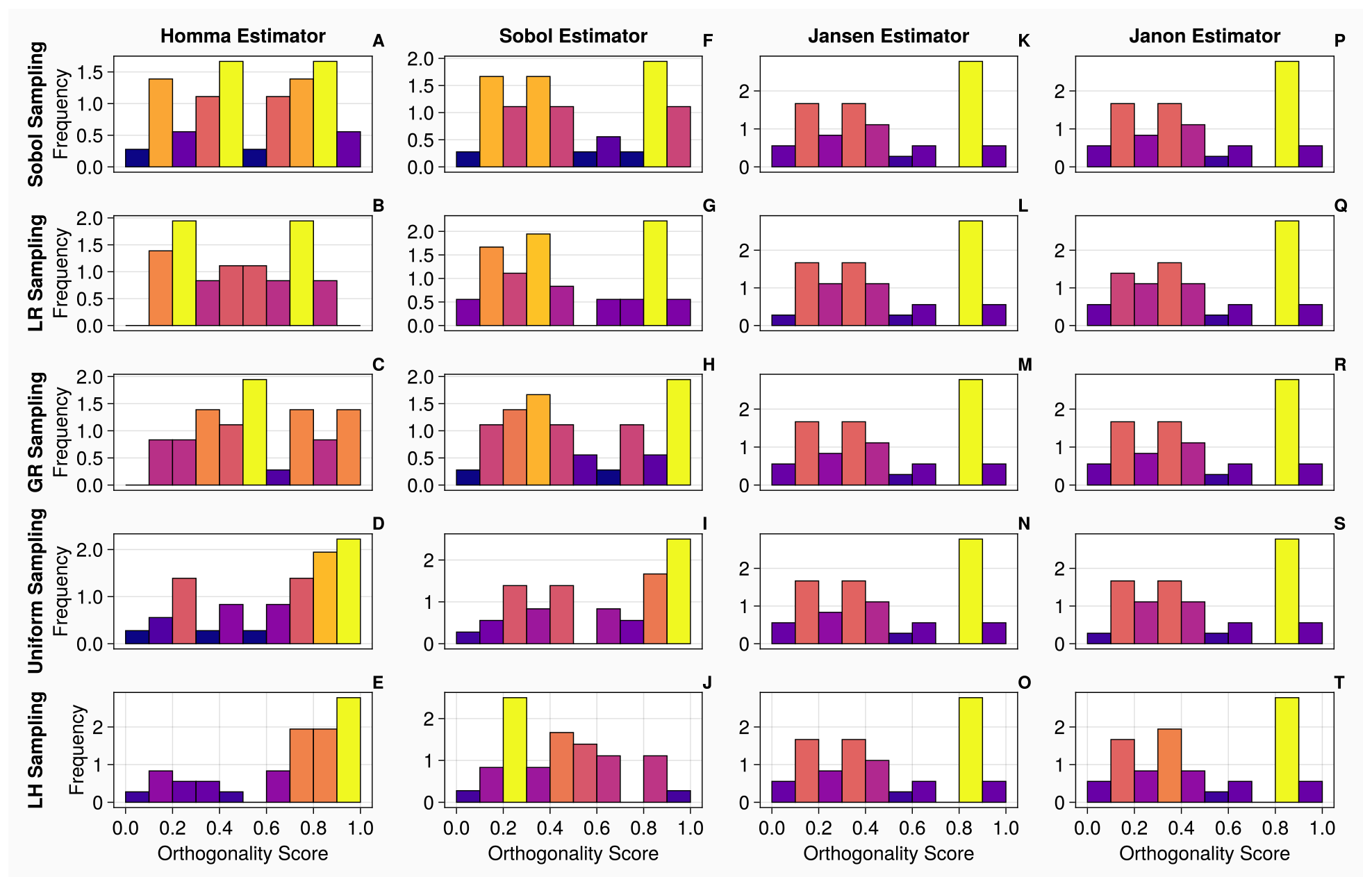
Orthogonality distributions of input parameters for the 1-chamber model with discrete measurements. Histograms A-T show the distribution of orthogonality returned from examinations of the sensitivity vectors, calculated from continuous measurements. Here, an orthogonality score of 1 represents total independence of input parameters, whereas 0 represents total dependence. Each individual diagram denotes a specific combination of sampling methodology and estimator type. The frequency of each histogram is normalised such that it is comparable between plots, i.e., the larger the frequency of a bin, the larger the number of orthogonality scores calculated from the original sensitivity vectors.

In Table 6, stratifying by estimator type and examining the range an input parameter exhibits across all sampling methodologies reveals that the Jansen and Janon estimators exhibit no variation for any input parameter set, given any sampling methodology, once again implying they are the optimum choice. The Homma and Sobol estimators exhibit variations of 5.11 to 2.22, respectively, upon the input parameter set. The variations for the discrete measurements are much greater than the variations seen with continuous measurements. When stratifying by sampling type, Table 7 shows the Sobol sampling method exhibits the smallest mean variation of an input set across all estimator types. As above, due to only the Sobol and Homma estimator exhibiting largely varying parameter rankings, stratifying by sampling methodology places more emphasis on the apparent less robust estimators. From this result, it does appear the Sobol sampling method may improve the robustness associated with a parameter orthogonality ranking. One notable observation as seen in Table 2 and Table 5 is that while the robustness of the Jansen and Janon estimator can be observed in both tables, the ranking associated with the orthogonality of input parameters changes quite dramatically (for example, *E*_*min*_, *R*_*mv*_, *Z*_*ao*_ and *R*_*s*_), highlighting how the change in data type may have consequences in parameter interpretation when conducting a sensitivity analysis.

**Table 6:** The range of parameter ranking across 5 sampling types for a specific estimator for the 1-chamber model with discrete measurements.

| | $\tau_{es}$ | $\tau_{ep}$ | $R_{mv}$ | $Z_{ao}$ | $R_s$ | $C_{sa}$ | $C_{sv}$ | $E_{max}$ | $E_{min}$ | Mean Variation of input set |
| --- | --- | --- | --- | --- | --- | --- | --- | --- | --- | --- |
| Range Homma | 3 | 3 | 4 | 4 | 6 | 6 | 7 | 5 | 8 | 5.11 |
| Range Sobol | 6 | 2 2/2024 | 1 | 1 | 3 | 3 | 1 | 2 | 1 | 2.22 |
| Range Jansen | 0 | 0 | 0 | 0 | 0 | 0 | 0 | 0 | 0 | 0 |
| Range Janon | 0 | 0 | 0 | 0 | 0 | 0 | 0 | 0 | 0 | 0 |

**Table 7:** The ranges of input parameters across 4 estimator types for a specific sampling method for the single ventricle model with discrete measurements.

| | $\tau_{es}$ | $\tau_{ep}$ | $R_{mv}$ | $Z_{ao}$ | $R_s$ | $C_{sa}$ | $C_{sv}$ | $E_{min}$ | $E_{max}$ | Mean variation of input set |
| --- | --- | --- | --- | --- | --- | --- | --- | --- | --- | --- |
| Range SS | 2 | 1 | 1 | 1 | 1 | 1 | 1 | 0 | 1 | 1.11 |
| Range LR | 3 | 1 | 1 | 1 | 3 | 2 | 2 | 3 | 3 | 2.11 |
| Range GR | 3 | 2 | 3 | 0 | 3 | 4 | 1 | 5 | 6 | 3.00 |
| Range U | 4 | 1 | 3 | 5 | 3 | 6 | 2 | 2 | 8 | 3.56 |
| Range LH | 4 | 3 | 2 | 5 | 3 | 3 | 6 | 2 | 1 | 3.22 |

Overall, for the 1-chamber model with 9 input parameters, we have seen consistent themes of the Jansen and Janon estimators being most robust and most reliable which appear attributable to the excellent convergence exhibited by these estimators. Sobol and Homma estimators exhibit very variable parameter rankings across different sampling methodologies which are in line with the poor convergence of these estimators. The Sobol sampling method appears to reduce the level of uncertainty associated with an input parameter’s orthogonality ranking, as shown in Tables 4 and 7 Interestingly, continuous measurements appear to reduce the level of variation associated with parameter orthogonality ranking when compared to discrete measurements.

### 3.3 2-chamber Model

In this section, we present the results of our uncertainty study on the 20 parameters, 2-chamber model. First, the uncertainty associated with the computation of Sobol total order indices, for this more complex model, with *N* = 20, 000 samples, is presented in Figure 8. Only the results for the left ventricular maximum elastance 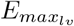 are displayed here. The Jansen and Janon estimators display identical index and confidence interval calculations, apart from when combined with the lattice rule sampling methodology, the plots show slightly larger confidence intervals for the left ventricular volume comparing to the other cases. The Homma and Sobol estimators again display much larger confidence interval estimates compared to the Jansen and Janon estimators. The errors associated with the Sobol sampling methodology when the Homma and Sobol estimator are used, are much smaller compared to the Latin hypercube and uniform sampling methodologies, hence demonstrating the impact sampling methodology can have on the estimations of sensitivity indices.

**Figure 8.**
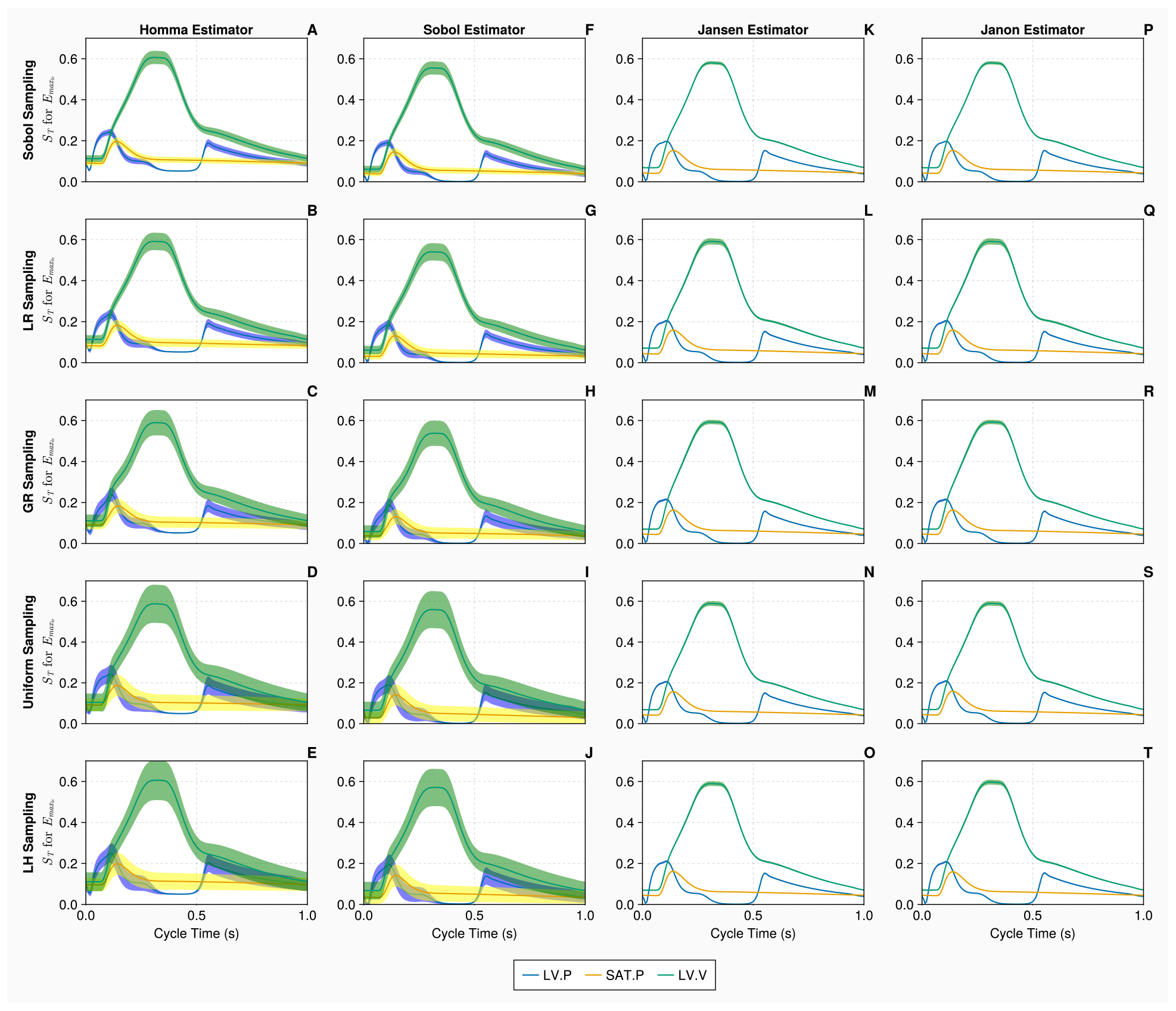
Total order Sobol indices *S*_*T*_ of the maximal left ventricular elastance 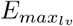 for the 2-chamber model with continuous measurements. Panels A - T show *S*_*T*_ of 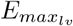, for 3 continuous measurements - left ventricular pressure, systemic arterial pressure and the left ventricular volume (represented in blue, yellow and green curves, respectively), over a single cardiac cycle with differing estimators and sampling methodologies. Measurements are evaluated with N = 20, 000 samples, using B = 1000 bootstrapped samples to evaluate the uncertainty of the estimate. The bands represent 95% confidence intervals associated with specific indices displayed as solid curves.

Next, we calculate total order indices on the 20 dimensional 2-chamber model with continuous measurements. From the histograms presented in Figure 9, the orthogonality spreads for the Jansen and Janon estimators are nearly identical for all sampling methodologies, excluding the uniform sampling, which shares the same orthogonality spread between the two estimators. With the Homma estimator, the spreads of orthogonality appear more consistent amongst the sampling techniques associated with itself, comparing against the test cases on the 1-chamber model, however, they are largely different from what returned from the Jansen and Janon estimators. The Sobol estimator generated results which appear closer to the orthogonality distributions of the Jansen and Janon estimators, whilst they are not identical, the orthogonality distributions between different sampling methodologies are more consistent than the ones exhibited by the Homma estimator. Examining Table 8, the rankings of input parameters are more consistent for the Jansen and Janon estimators, although there are slight discrepancies, as seen on the simple 1-chamber model.

**Table 8:** Input parameter ranking for the 2-chamber model with continuous measurements. Here, input parameters are ranked based on the averaged orthogonality score returned from the calculated total order sensitivity matrix. In addition, the ranking is stratified by both sampling and estimator types.

|  | Homma |  |  |  |  | Sobol |  |  |  |  | Jansen |  |  |  |  | Janon |  |  |  |  |
| --- | --- | --- | --- | --- | --- | --- | --- | --- | --- | --- | --- | --- | --- | --- | --- | --- | --- | --- | --- | --- |
|  | SS | LR | GR | U | LH | SS | LR | GR | U | LH | SS | LR | GR | U | LH | SS | LR | GR | U | LH |
| $E_{min_{lv}}$ | 3 | 8 | 8 | 2 | 2 | 12 | 8 | 13 | 9 | 11 | 5 | 5 | 5 | 6 | 5 | 5 | 5 | 6 | 6 | 5 |
| $E_{max_{lv}}$ | 7 | 11 | 15 | 1 | 1 | 2 | 3 | 3 | 5 | 5 | 1 | 1 | 1 | 1 | 1 | 1 | 1 | 1 | 1 | 1 |
| $\tau_{es_{lv}}$ | 1 | 4 | 6 | 6 | 8 | 8 | 5 | 8 | 8 | 8 | 3 | 3 | 3 | 4 | 3 | 3 | 3 | 3 | 4 | 3 |
| $\tau_{ep_{lv}}$ | 2 | 5 | 7 | 7 | 9 | 9 | 4 | 6 | 6 | 9 | 2 | 2 | 2 | 2 | 2 | 2 | 2 | 2 | 2 | 2 |
| $E_{min_{la}}$ | 8 | 10 | 1 | 12 | 11 | 13 | 14 | 7 | 2 | 14 | 7 | 7 | 7 | 7 | 7 | 7 | 7 | 7 | 7 | 7 |
| $E_{max_{la}}$ | 9 | 9 | 2 | 11 | 12 | 16 | 17 | 18 | 14 | 15 | 18 | 18 | 18 | 19 | 18 | 18 | 18 | 18 | 19 | 18 |
| $\tau_{es_{la}}$ | 19 | 16 | 16 | 18 | 19 | 19 | 19 | 15 | 20 | 13 | 8 | 8 | 8 | 16 | 9 | 8 | 8 | 8 | 16 | 13 |
| $\tau_{ep_{la}}$ | 13 | 1 | 11 | 13 | 17 | 15 | 15 | 20 | 18 | 20 | 16 | 16 | 17 | 17 | 16 | 16 | 16 | 17 | 17 | 15 |
| $Z_{ao}$ | 10 | 2 | 13 | 14 | 13 | 11 | 10 | 12 | 7 | 10 | 4 | 4 | 4 | 3 | 4 | 4 | 4 | 4 | 3 | 4 |
| $R_{mv}$ | 17 | 15 | 12 | 3 | 5 | 18 | 16 | 17 | 10 | 18 | 17 | 17 | 16 | 15 | 17 | 17 | 17 | 16 | 15 | 17 |
| $C_{sas}$ | 16 | 17 | 18 | 20 | 20 | 7 | 6 | 4 | 17 | 16 | 9 | 9 | 9 | 8 | 8 | 9 | 9 | 9 | 8 | 8 |
| $R_{sas}$ | 18 | 18 | 19 | 16 | 16 | 14 | 7 | 14 | 4 | 12 | 6 | 6 | 6 | 5 | 6 | 6 | 6 | 5 | 5 | 6 |
| $L_{sas}$ | 14 | 20 | 17 | 17 | 15 | 1 | 2 | 2 | 1 | 1 | 15 | 15 | 15 | 14 | 15 | 15 | 15 | 15 | 14 | 16 |
| $C_{sat}$ | 6 | 3 | 3 | 4 | 4 | 6 | 11 | 9 | 12 | 7 | 10 | 10 | 10 | 9 | 10 | 10 | 10 | 10 | 9 | 9 |
| $R_{sat}$ | 12 | 12 | 14 | 15 | 18 | 3 | 9 | 5 | 19 | 6 | 13 | 11 | 12 | 11 | 13 | 13 | 11 | 12 | 10 | 10 |
| $L_{sat}$ | 15 | 19 | 20 | 19 | 14 | 10 | 1 | 1 | 3 | 2 | 14 | 14 | 14 | 13 | 14 | 14 | 14 | 14 | 13 | 14 |
| $R_{sar}$ | 5 | 6 | 4 | 9 | 6 | 5 | 12 | 11 | 16 | 3 | 12 | 13 | 13 | 12 | 12 | 12 | 13 | 13 | 12 | 12 |
| $R_{scp}$ | 4 | 7 | 5 | 10 | 7 | 4 | 13 | 10 | 15 | 4 | 11 | 12 | 11 | 10 | 11 | 11 | 12 | 11 | 11 | 12 |
| $R_{svn}$ | 20 | 14 | 9 | 5 | 10 | 20 | 18 | 16 | 11 | 19 | 19 | 19 | 19 | 18 | 19 | 19 | 19 | 19 | 18 | 19 |
| $C_{svn}$ | 11 | 13 | 10 | 8 | 3 | 17 | 20 | 19 | 13 | 17 | 20 | 20 | 20 | 20 | 20 | 20 | 20 | 20 | 20 | 20 |

**Figure 9.**
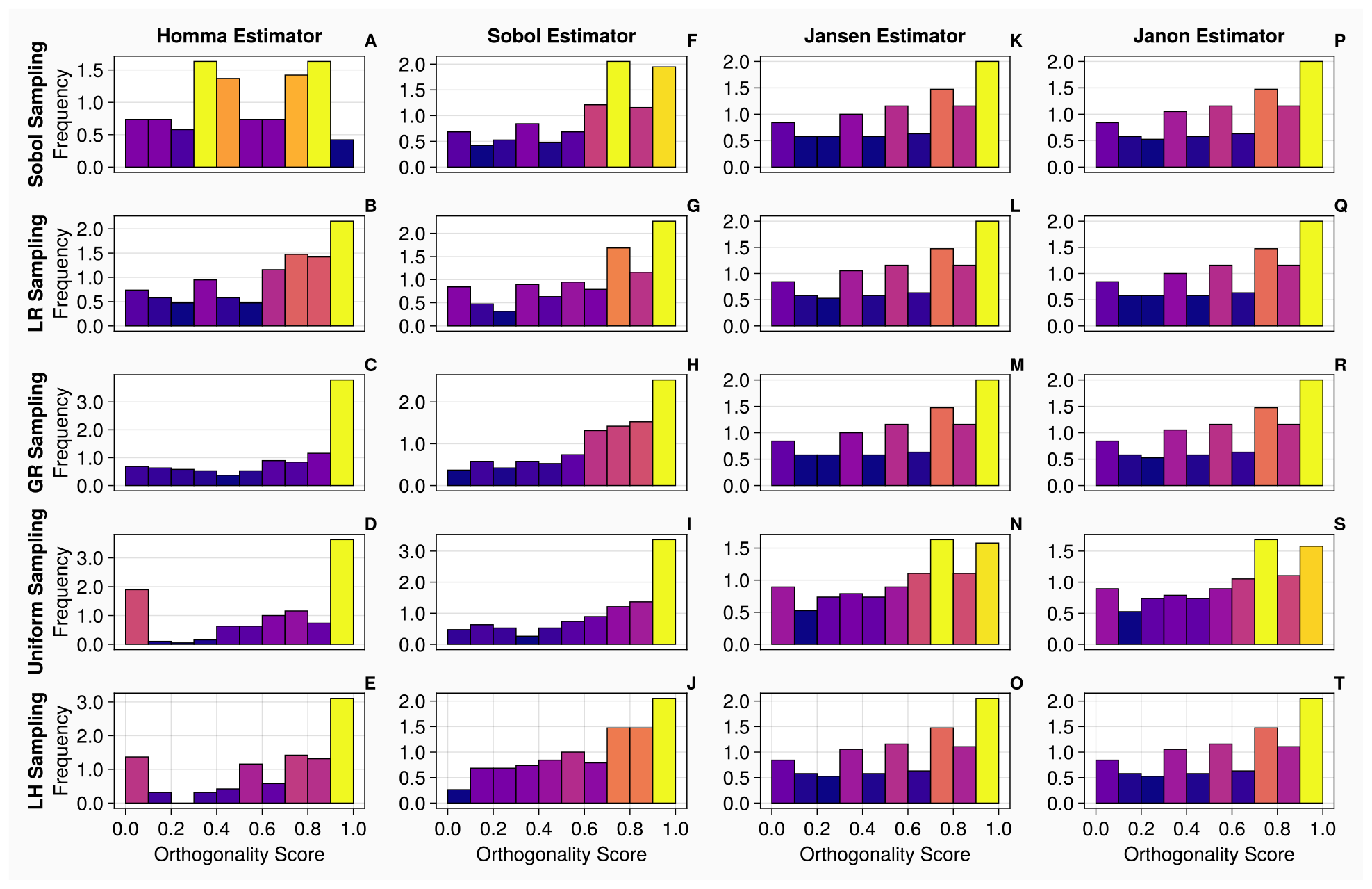
Orthogonality distributions of input parameters for the 2-chamber model with continuous measurements. Histograms A-T show the distribution of orthogonality returned from examinations of the sensitivity vectors, calculated from continuous measurements. Here, an orthogonality score of 1 represents total independence of input parameters, whereas 0 represents total dependence. Each individual diagram denotes a specific combination of sampling methodology and estimator type. The frequency of each histogram is normalised such that it is comparable between plots, i.e., the larger the frequency of a bin, the larger the number of orthogonality scores calculated from the original sensitivity vectors.

In Table 9, stratifying by estimator type and examining the range an input parameter exhibits across all sampling methodologies reveal that the Jansen and Janon estimators exhibit minimal variations to sampling methodologies - 1.3 and 1.45, respectively. The Homma and Sobol estimators exhibit variations of 8.2 and 7.35 respectively upon the input parameter set. When stratifying by sampling type, Table 10 shows the Sobol sampling method exhibits the smallest mean variation of an input set across all estimator types of 7.2. This is still a large variation due to all estimator types being considered and therefore the range accounts for some of the spurious values generated by the Homma and Sobol estimators.

**Table 9:** The ranges of input parameters across 5 sampling types for a specific estimator for the 2-chamber model with continuous measurements.

|  | Range Homma | Range Sobol | Range Jansen | Range Janon |
| --- | --- | --- | --- | --- |
| $E_{min_{lv}}$ | 6 | 5 | 1 | 1 |
| $E_{max_{lv}}$ | 14 | 3 | 0 | 0 |
| $\tau_{es_{lv}}$ | 7 | 3 | 1 | 1 |
| $\tau_{ep_{lv}}$ | 7 | 5 | 0 | 0 |
| $E_{min_{la}}$ | 11 | 12 | 0 | 0 |
| $E_{max_{la}}$ | 10 | 4 | 1 | 1 |
| $\tau_{es_{la}}$ | 3 | 7 | 8 | 8 |
| $\tau_{ep_{la}}$ | 16 | 5 | 1 | 2 |
| $Z_{ao}$ | 12 | 5 | 1 | 1 |
| $R_{mv}$ | 14 | 8 | 2 | 2 |
| $C_{sas}$ | 4 | 13 | 1 | 1 |
| $R_{sas}$ | 3 | 7 | 1 | 1 |
| $L_{sas}$ | 6 | 1 | 1 | 2 |
| $C_{sat}$ | 3 | 6 | 1 | 1 |
| $R_{sat}$ | 6 | 16 | 2 | 3 |
| $L_{sat}$ | 6 | 9 | 1 | 1 |
| $R_{sar}$ | 5 | 11 | 1 | 2 |
| $R_{scp}$ | 6 | 11 | 2 | 1 |
| $R_{svn}$ | 15 | 9 | 1 | 1 |
| $C_{svn}$ | 10 | 7 | 0 | 0 |
| Mean Variation Of Input Set | 8.2 | 7.35 | 1.3 | 1.45 |

**Table 10:** The ranges of input parameters across 4 estimator types for a specific sampling method for the 2-chamber model with continuous measurements.

|  | Range SS | Range LR | Range GR | Range U | Range LH |
| --- | --- | --- | --- | --- | --- |
| $E_{minlv}$ | 9 | 3 | 7 | 7 | 9 |
| $E_{maxlv}$ | 6 | 10 | 14 | 4 | 4 |
| $\tau_{eslv}$ | 7 | 2 | 5 | 4 | 5 |
| $\tau_{eplv}$ | 7 | 3 | 5 | 5 | 7 |
| $E_{minla}$ | 6 | 7 | 6 | 10 | 7 |
| $E_{maxla}$ | 9 | 9 | 16 | 9 | 6 |
| $\tau_{esla}$ | 11 | 11 | 8 | 4 | 10 |
| $\tau_{epla}$ | 3 | 15 | 4 | 5 | 5 |
| $Z_{ao}$ | 7 | 8 | 8 | 11 | 9 |
| $R_{mv}$ | 1 | 2 | 5 | 12 | 13 |
| $C_{sas}$ | 9 | 11 | 14 | 12 | 12 |
| $R_{sas}$ | 12 | 12 | 14 | 11 | 10 |
| $L_{sas}$ | 14 | 18 | 15 | 16 | 15 |
| $C_{sat}$ | 4 | 8 | 7 | 8 | 6 |
| $R_{sat}$ | 10 | 3 | 9 | 9 | 12 |
| $L_{sat}$ | 5 | 18 | 13 | 16 | 12 |
| $R_{sar}$ | 7 | 7 | 9 | 7 | 9 |
| $R_{scp}$ | 7 | 4 | 7 | 5 | 8 |
| $R_{svn}$ | 1 | 5 | 10 | 13 | 9 |
| $C_{svn}$ | 9 | 7 | 10 | 12 | 17 |
| Mean Variation<br>Of Input Set | 7.2 | 8.15 | 9.3 | 10.65 | 9.75 |

Figure 10 displays the convergence and uncertainty associated with the computation of the total order indices of the venous compliance *C*_*svn*_, for the 2-chamber model, against the discrete measurements. We see in all cases that as the sample size is increased, the estimate and uncertainty associated with the indices improve. Similar as the continuous measurement case for 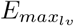 shown in Fig 8, the Jansen and Janon estimators provide the most efficient convergence and the smallest error when calculating the indices. A sample size of *N* = 20, 000 is taken for the discrete measurements, as seen in the Jansen and Janon columns (Panels K - T), any additional sampling would return minimal improvements in terms of accurate calculation of the indices. The Homma and Sobol estimators display errors which are considerably larger than that of Jansen and Janon estimators. We see when the upper limit sample size of *N* = 30, 000 is reached, the Homma and Sobol estimator errors are still large. This results demonstrates that for this complex model, less efficient estimators (such as Homma and Sobol) and a less accurate sampling method (such as Latin hypercube) display large confidence intervals and struggle to return converged index values.

**Figure 10.**
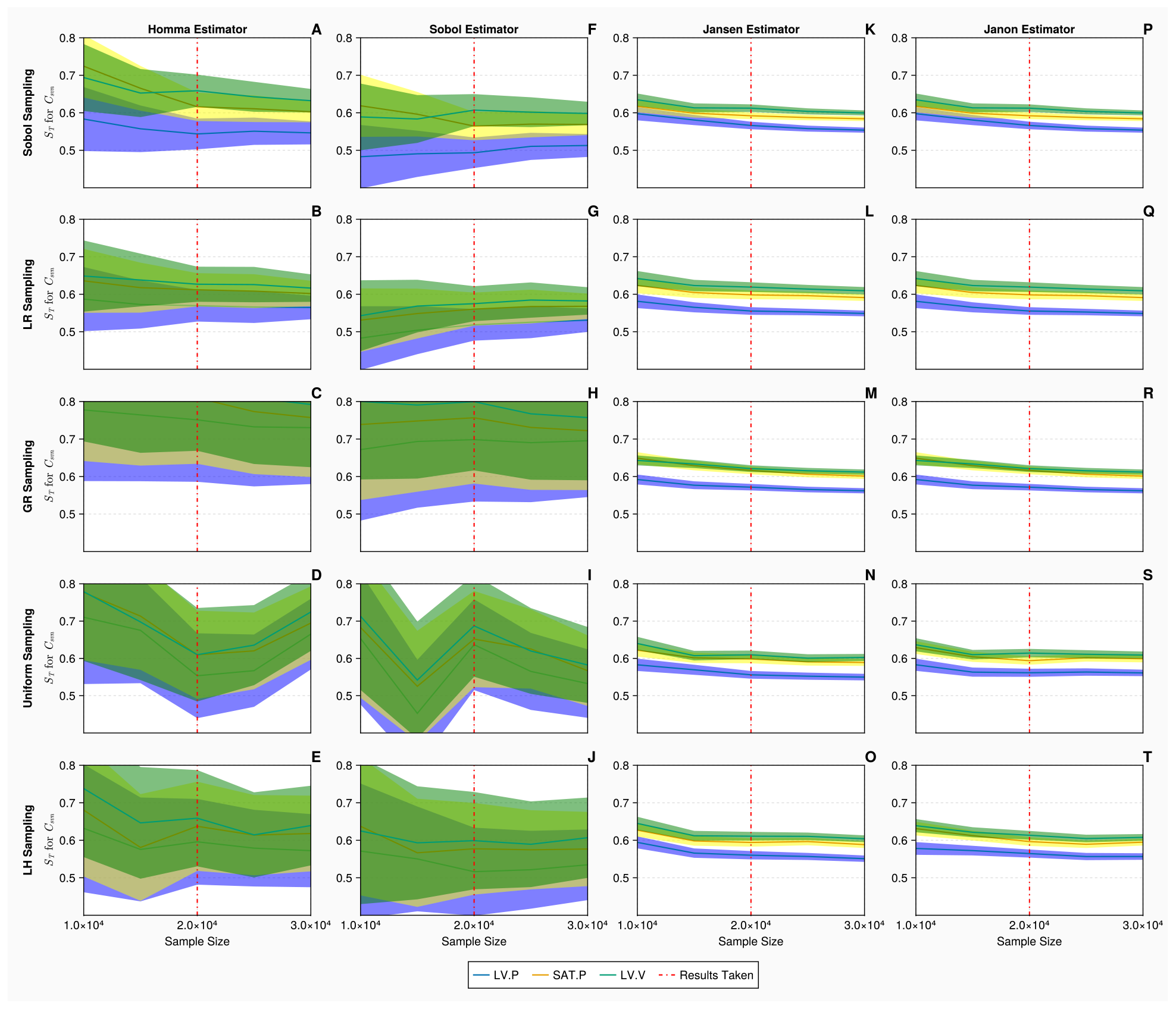
Total order Sobol indices *S*_*T*_ of the mitral valve resistance *C*_*svn*_ for the 2-chamber model with discrete measurements. Panels A - T show *S*_*T*_ of *C*_*svn*_, for 3 discrete measurements: mean left ventricular pressure, maximum systemic arterial pressure and maximum left ventricular volume (represented in blue, yellow and green, respectively), evaluated at increasing sample sizes (*N*∈ [10000, 30000] using *B* = 1000 bootstrapped samples), with differing estimators and sampling methodologies. The bands represent 95% confidence intervals associated with specific indices displayed as solid curves. The red solid vertical lines represent the point (*N* = 20, 000) at which the sample size is taken.

From the histograms presented in Figure 11, the orthogonality spreads exhibit similar trends to that of the continuous measurements, shown in Figure 9. We note that the histograms are identical for the Jansen and Janon except when the Uniform sampling method is used (which exhibits slight variations from the other histograms). With the Homma and Sobol estimators, although there appears to be low level consistency amongst their orthogonality distributions, they are very different to the ones produced by the Jansen and Janon estimators. Examining Table 11, the rankings of input parameters are consistent for the Jansen and Janon estimators apart from the Uniform column which often returns a parameter ranking differing to the other sampling types.

**Table 11:** Input parameter ranking for the 2-chamber model with discrete measurements. Here, input parameters are ranked based on the averaged orthogonality score returned from the calculated total order sensitivity matrix. In addition, the ranking is stratified by both sampling and estimator types.

|  | Homma |  |  |  |  | Sobol |  |  |  |  | Jansen |  |  |  |  | Janon |  |  |  |  |
| --- | --- | --- | --- | --- | --- | --- | --- | --- | --- | --- | --- | --- | --- | --- | --- | --- | --- | --- | --- | --- |
|  | SS | LR | GR | U | LH | SS | LR | GR | U | LH | SS | LR | GR | U | LH | SS | LR | GR | U | LH |
| $E_{min_{lv}}$ | 17 | 18 | 14 | 1 | 3 | 7 | 6 | 6 | 9 | 6 | 5 | 5 | 5 | 5 | 5 | 5 | 5 | 5 | 5 | 5 |
| $E_{max_{lv}}$ | 18 | 14 | 9 | 3 | 7 | 19 | 18 | 19 | 20 | 20 | 20 | 20 | 20 | 19 | 20 | 20 | 20 | 20 | 19 | 20 |
| $\tau_{es_{lv}}$ | 4 | 5 | 4 | 10 | 11 | 8 | 7 | 8 | 6 | 5 | 4 | 4 | 4 | 4 | 4 | 4 | 4 | 4 | 4 | 4 |
| $\tau_{ep_{lv}}$ | 1 | 1 | 1 | 2 | 8 | 4 | 4 | 7 | 7 | 4 | 3 | 3 | 3 | 3 | 3 | 3 | 3 | 3 | 3 | 3 |
| $E_{min_{la}}$ | 19 | 19 | 8 | 14 | 12 | 14 | 14 | 14 | 1 | 16 | 16 | 16 | 16 | 18 | 16 | 16 | 16 | 16 | 18 | 16 |
| $E_{max_{la}}$ | 20 | 20 | 6 | 9 | 9 | 15 | 13 | 16 | 15 | 9 | 17 | 17 | 17 | 16 | 17 | 17 | 17 | 17 | 16 | 17 |
| $\tau_{es_{la}}$ | 9 | 12 | 17 | 19 | 17 | 16 | 19 | 18 | 17 | 15 | 12 | 12 | 12 | 12 | 12 | 12 | 12 | 12 | 12 | 12 |
| $\tau_{ep_{la}}$ | 15 | 17 | 7 | 12 | 15 | 17 | 17 | 17 | 18 | 7 | 15 | 15 | 15 | 17 | 15 | 15 | 15 | 15 | 17 | 15 |
| $Z_{ao}$ | 14 | 6 | 12 | 20 | 13 | 3 | 2 | 5 | 4 | 2 | 1 | 1 | 1 | 1 | 1 | 1 | 1 | 1 | 1 | 1 |
| $R_{mv}$ | 13 | 15 | 10 | 4 | 5 | 18 | 15 | 15 | 16 | 17 | 14 | 14 | 14 | 15 | 14 | 14 | 14 | 14 | 15 | 14 |
| $C_{sas}$ | 11 | 7 | 16 | 15 | 16 | 12 | 16 | 11 | 2 | 18 | 7 | 7 | 7 | 7 | 7 | 7 | 7 | 7 | 7 | 7 |
| $R_{sas}$ | 7 | 10 | 19 | 18 | 20 | 6 | 3 | 4 | 5 | 3 | 2 | 2 | 2 | 2 | 2 | 2 | 2 | 2 | 2 | 2 |
| $L_{sas}$ | 8 | 8 | 18 | 17 | 19 | 1 | 1 | 1 | 3 | 1 | 18 | 18 | 18 | 14 | 18 | 18 | 18 | 18 | 14 | 18 |
| $C_{sat}$ | 5 | 4 | 5 | 11 | 6 | 5 | 5 | 3 | 12 | 12 | 6 | 6 | 6 | 6 | 6 | 6 | 6 | 6 | 6 | 6 |
| $R_{sat}$ | 10 | 11 | 15 | 13 | 14 | 11 | 11 | 2 | 8 | 8 | 10 | 11 | 9 | 9 | 10 | 10 | 11 | 9 | 9 | 10 |
| $L_{sat}$ | 6 | 9 | 20 | 16 | 18 | 2 | 10 | 12 | 13 | 13 | 8 | 8 | 8 | 8 | 8 | 8 | 8 | 8 | 8 | 8 |
| $R_{sar}$ | 3 | 2 | 2 | 8 | 4 | 9 | 8 | 9 | 11 | 11 | 9 | 9 | 10 | 11 | 11 | 9 | 9 | 10 | 11 | 11 |
| $R_{scp}$ | 2 | 3 | 3 | 7 | 2 | 10 | 9 | 10 | 10 | 10 | 11 | 10 | 11 | 10 | 9 | 11 | 10 | 11 | 10 | 9 |
| $R_{svn}$ | 16 | 16 | 13 | 6 | 1 | 13 | 12 | 13 | 14 | 14 | 13 | 13 | 13 | 13 | 13 | 13 | 13 | 13 | 13 | 13 |
| $C_{svn}$ | 12 | 13 | 11 | 5 | 10 | 20 | 20 | 20 | 19 | 19 | 19 | 19 | 19 | 20 | 19 | 19 | 19 | 19 | 20 | 19 |

**Figure 11.**
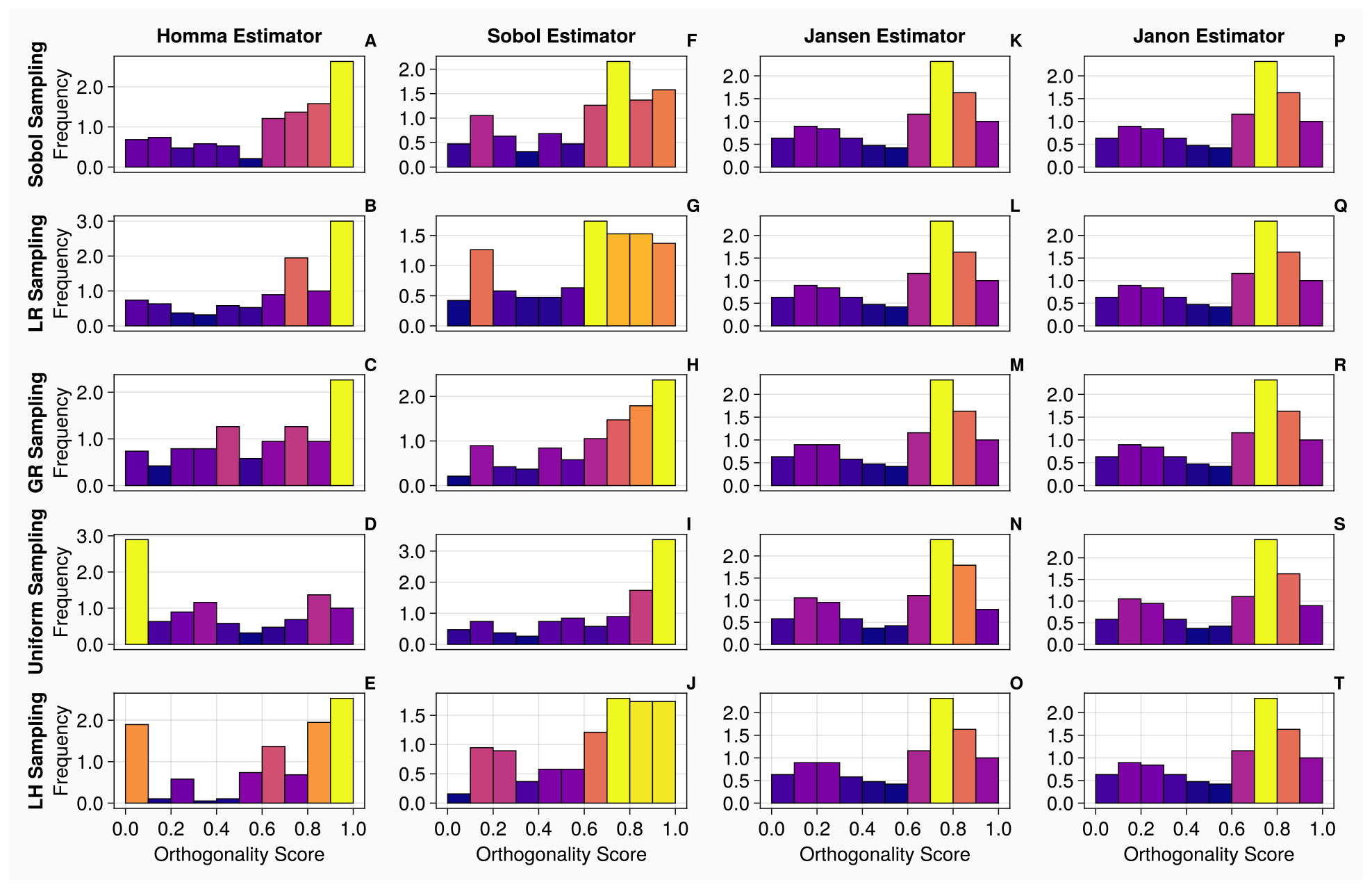
Orthogonality distributions of input parameters for the 2-chamber model with discrete measurements. Histograms A-T show the distribution of orthogonality returned from examinations of the sensitivity vectors, calculated from continuous measurements. Here, an orthogonality score of 1 represents total independence of input parameters, whereas 0 represents total dependence. Each individual diagram denotes a specific combination of sampling methodology and estimator type. The frequency of each histogram is normalised such that it is comparable between plots, i.e., the larger the frequency of a bin, the larger the number of orthogonality scores calculated from the original sensitivity vectors.

**Figure 12.**
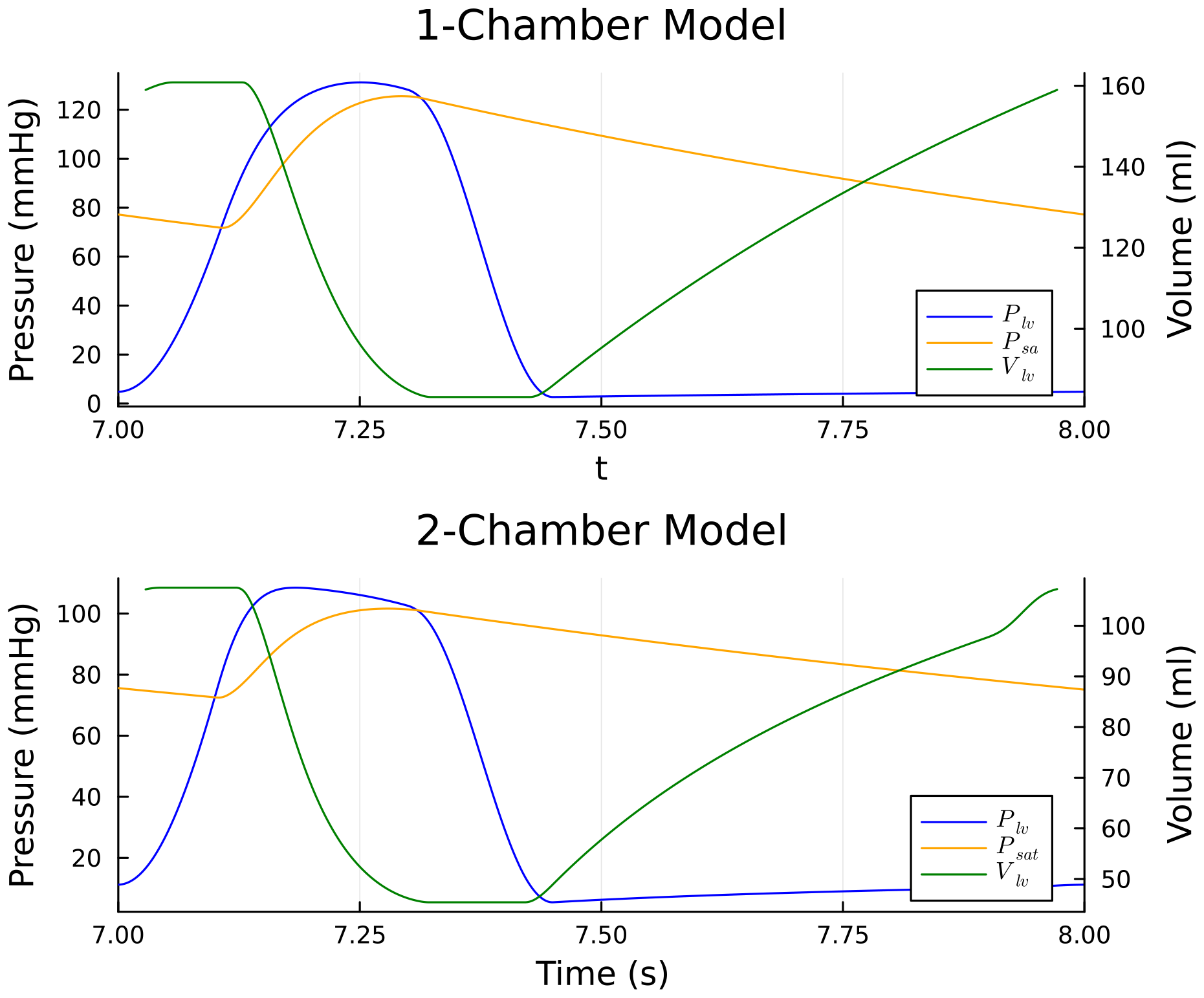
Time Series solutions for the 1 and 2 chamber cardiovascular models investigated in this work. The solutions shown are the ones which are utilised in the investigation.

In Table 12, stratifying by estimator type and examining the range a input parameter exhibits across all sampling methodologies reveals that the Jansen and Janon estimators exhibit minimal variation to sampling methodologies - 0.9 and 1.0, respectively. This is an improvement on the continuous measurements as the Jansen estimator returns less than 1 parameter range variation. The Homma and Sobol estimators produce variations of 10.9 and 5.55 respectively upon the input parameter set. When stratifying by sampling type, Table 13 shows the Lattice Rule sampling method exhibits the smallest mean variation of an input set across all estimator types of 5.75. This is still a large variation but is less than the variation exhibited by the continuous measurements.

**Table 12:** The ranges of input parameters across 5 sampling types for a specific estimator for the systemic circulation model with discrete measurements.

|  | Range Homma | Range Sobol | Range Jansen | Range Janon |
| --- | --- | --- | --- | --- |
| $E_{min_{lv}}$ | 17 | 3 | 0 | 0 |
| $E_{max_{lv}}$ | 15 | 2 | 1 | 1 |
| $\tau_{es_{lv}}$ | 7 | 3 | 0 | 0 |
| $\tau_{ep_{lv}}$ | 6 | 3 | 0 | 2 |
| $E_{min_{la}}$ | 11 | 15 | 2 | 1 |
| $E_{max_{la}}$ | 14 | 7 | 1 | 0 |
| $\tau_{es_{la}}$ | 10 | 4 | 0 | 2 |
| $\tau_{ep_{la}}$ | 10 | 11 | 2 | 2 |
| $Z_{ao}$ | 14 | 3 | 0 | 0 |
| $R_{mv}$ | 11 | 3 | 1 | 1 |
| $C_{sas}$ | 9 | 16 | 0 | 0 |
| $R_{sas}$ | 13 | 3 | 0 | 0 |
| $L_{sas}$ | 11 | 2 | 4 | 4 |
| $C_{sat}$ | 7 | 9 | 0 | 0 |
| $R_{sat}$ | 5 | 9 | 2 | 2 |
| $L_{sat}$ | 14 | 11 | 0 | 0 |
| $R_{sar}$ | 6 | 3 | 2 | 2 |
| $R_{scp}$ | 5 | 1 | 2 | 2 |
| $R_{svn}$ | 15 | 2 | 0 | 0 |
| $C_{svn}$ | 7 | 1 | 1 | 1 |
| Mean Variation Of Input Set | 10.9 | 5.55 | 0.9 | 1.0 |

**Table 13:** The ranges of input parameters across 4 estimator types for a specific sampling method for the 2-chamber model with discrete measurements.

|  | Range SS | Range LR | Range GR | Range U | Range LH |
| --- | --- | --- | --- | --- | --- |
| $E_{minlv}$ | 12 | 13 | 9 | 8 | 3 |
| $E_{maxlv}$ | 2 | 6 | 11 | 17 | 13 |
| $\tau_{eslv}$ | 5 | 3 | 4 | 6 | 7 |
| $\tau_{eplv}$ | 3 | 3 | 6 | 5 | 5 |
| $E_{minla}$ | 5 | 5 | 8 | 17 | 4 |
| $E_{maxla}$ | 5 | 7 | 11 | 7 | 8 |
| $\tau_{esla}$ | 7 | 7 | 6 | 7 | 5 |
| $\tau_{epla}$ | 2 | 2 | 10 | 6 | 8 |
| $Z_{ao}$ | 13 | 5 | 11 | 19 | 12 |
| $R_{mv}$ | 5 | 1 | 5 | 12 | 12 |
| $C_{sas}$ | 5 | 9 | 9 | 13 | 11 |
| $R_{sas}$ | 5 | 8 | 17 | 16 | 18 |
| $L_{sas}$ | 17 | 17 | 17 | 14 | 18 |
| $C_{sat}$ | 1 | 2 | 3 | 6 | 6 |
| $R_{sat}$ | 1 | 0 | 13 | 5 | 6 |
| $L_{sat}$ | 6 | 2 | 12 | 8 | 10 |
| $R_{sar}$ | 6 | 7 | 8 | 3 | 7 |
| $R_{scp}$ | 9 | 7 | 8 | 3 | 7 |
| $R_{svn}$ | 3 | 4 | 0 | 7 | 13 |
| $C_{svn}$ | 8 | 7 | 9 | 15 | 9 |
| Mean Variation<br>Of Input Set | 6 | 5.75 | 9.35 | 9.7 | 9.15 |

Overall, for the more complex 2-chamber model with 20 input parameters, the Jansen and Janon estimators are consistently the most robust and reliable estimators. When using continuous measurements, neither returns an input parameter set mean variation greater than 1. When using discrete measurements, they return a mean variation less than or equal to 1 (see Tables 9 and 12). This, as in the 1-chamber case, could be attributable to the efficient rate of convergence displayed by the Jansen and Janon estimator. The Sobol and Homma estimators exhibit very different parameter rankings across different sampling methodologies with variations of up to 10.9. These large variations are in line with the poor convergence associated with these estimators. The Sobol and Lattice Rule sampling method appears to reduce the level of uncertainty associated with an input parameter’s orthogonality ranking (see Tables 10 and 13), despite spurious parameter rankings from the Homma and Sobol estimators leading to large parameter variation when stratified by sampling methodologies.

## 4 Discussion

Utilising two cardiovascular system models, the main aim of our investigation is to test the robustness of the calculation of the input parameter orthogonality, while varying total order estimator types and sampling methodologies, across differing input parameter dimensionalities and types of data on which the total order indices were calculated. The results presented in Section 3 display overwhelming robustness for the Jansen and Janon estimators when calculating total order indices when compared to other options. For the 1-chamber, 9-parameter model, we observed that these two estimators gave nearly invariant outcomes to both the sampling methodology and the data type. When the dimensionality of the model parameters is increased to 20, we noted that the Jansen and Janon estimators exhibited small variations on the input parameter orthogonality rankings. For the Jansen estimator with discrete measurements, it returned a mean variation of less than 1. We observed that the Homma and Sobol estimators regularly returned mean variations for input parameter sets greater than 1, which is particularly amplified when the model dimensionality is increased.

Given our aim is to assess the use-ability of estimators and sampling methodologies for practical identifiability studies, these results indicate that if the used estimator and sampling methodology are not robust, the calculated optimal parameter set is unreliable. The usage of unrobust estimators and sampling methods can therefore produce misleading conclusions with practical consequences, especially in investigations applied to life sciences. Interestingly, we witnessed that the variations attached to the Sobol and Lattice Rule sampling methods were the lowest across all model dimensionalities and data types. Our results also reinforce the findings reported in [14, 57] that the commonly used Latin Hypercube method is less than optimal in exploring the input parameter space, especially at high dimensionalites.

The Jansen and Janon robustness at calculating total order indices can be attributed to the Jansen estimator never allowing negative values in the numerator (see estimator definitions in Table 1), where as the Janon estimator is the only estimator which has been proven to be both asymptotically normally distributed and asymptotically efficient meaning as the sample size increases the estimation error associated with calculating the indices is negligible [47]. They are both highly optimised estimators with very little room for improvement [46, 47]. The allowance of negative indices in both the Sobol and Homma estimators is an explanation for the poor performance in the calculations of these indices.

With the highly optimised Jansen and Janon estimators, it is evident that they consistently exhibit the most efficient convergence and produce the smallest uncertainties when calculating total order indices. This phenomenon matches the consistent orthogonality observed among input parameters across various sampling techniques when coupled with the Jansen and Janon estimator. Conversely, the Homma and Sobol estimators tend to yield significantly larger uncertainties when sample sizes are held constant among estimators, thus explaining the lack of consistent orthogonality rankings for the input parameters. Increasing the sample size seems to ameliorate the uncertainties associated with the Homma and Sobol estimators, particularly when employing the Sobol, lattice rule, and Golden sampling methods. This observation underscores the resilience of low-discrepancy sequences, demonstrating their effectiveness even in conjunction with a sub-optimal estimator. While the work of Puy et al. [18] did not delve into extensive convergence or uncertainty quantification, it is plausible to infer that the Jansen and Janon estimators, with their superior convergence rates and lower uncertainty, played a pivotal role in their conclusion that these estimators are the most efficient at capturing the true effects of input parameters.

Our investigation has been confined to two highly non-linear stiff differential algebraic equation systems with the understanding that they represent a high level of complexity, therefore good modelling guidelines obtained here would be readily applicable to simpler, more linear models which are associated with a less variable input parameter space. Thus, obtaining sensitivity estimates for linear models is considerably less expensive than what has been conducted here. Convergence and uncertainty quantification have historically been left out of sensitivity analysis studies, despite being highlighted as vital, if the results of studies were then to influence policy /societal /clinical decisions [14, 58]. In this study, we have highlighted the impact that convergence has on a total order estimator, alongside this, we have also shown the impact of the level of sampling taken, on the calculation of total order estimators. It appears intuitively sensible that the higher density of sampling leads to better resolution of the input parameter space, hence our sensitivity analysis gives a better indication about which input parameters are truly influential. Current literature states that *N >* 500 but this recommendation is based on physical systems which are mostly linear. The work conducted and results shown in this study highlight that for a highly non-linear system, one should investigate *N >* 5000. More importantly, it is clear that no two systems are the same, so for one to ensure adequate resolution of an input parameter space, convergence and uncertainty quantification through bootstrapping must be an essential step in any modeller’s workflow, given the aim is to perform accurate and robust parameter identification studies.

Although not exhaustive, the list of estimators investigated in this paper does represent what are readily available, practically usable and computationally feasible. Puy et al. [18] also recommended the estimator introduced by Azzini et al. [59], which appeared to produce similar results to the Jansen estimator. The Azzini estimator requires *k* = 2*N* (*p* + 1) model evaluations, compared to *N* (*p* + 2) evaluations needed by the estimators investigated in this work. The 2-chamber model would require *k* = 1, 320, 000 model evaluations for Azzini estimator. For models with higher parameter dimensions, this would be computationally infeasible. With the increasing prospect of digital twins in healthcare, more complex and detailed models which accurately represent the true physiological processes are generated. However, the bottleneck which prevents the progression of these models to clinically applicable situations is the computational cost associated with a detailed sensitivity analysis. This does not refer to the calculation of the indices, but the process of solving the dynamical system. Therefore, while new estimators may prove to be accurate, the focus must be on efficient resolution of complex dynamical systems and the efficiency of the estimators for low sample numbers, in order to ensure a thorough sensitivity analysis.

All the estimators used in this work are available in the global sensitivity packages such as SALib, GlobalSensitivity.jl, SenSobol and sbiosobol [60, 54, 61]. It is reassuring to see that the default estimator used to calculate the total order index, in the available packages, is the Jansen estimator. Given the conclusion drawn from Puy et al. [18] and the findings from this work, researchers could straightforwardly use the above packages when performing practical identifiability studies and would obtain a reliable optimal set of input parameters which best describes the experimental data available to them.

Another area of research is the calculation of total order sensitivity indices where one assumes dependency between input parameters. This is partially investigated by Puy et al. [61] in implementing the method of Glen et al. [62]. This method requires a prescription of linear dependencies between parameters, however, these are often not known in realistic life science models. As a result, Puy et. al. demonstrated the inaccuracy associated with this method when calculating the true effects. There have been various methods deriving variance based sensitivity indices with dependent inputs [63, 64, 65], however, similar as the method of Glen et al. [62], they require knowledge of the dependencies that exist within the model and therefore the computational power needed to simulate these indices is often much larger than the standard Sobol indices. On top of this, there is no accepted method for how to calculate these dependent indices which should be of interest for future work. Therefore, the need to understand how input parameter orthogonality is affected by varying estimators and sampling methodologies is of significant importance, in order for total order sensitivity indices to be utilised in identifiability studies.

While we have conducted this work through the lens of utilising the method of Li et. al. [7] (see Eq. (1)) for practical identifiability studies, it is also applicable to other methods. Another approach of identifying input parameters is the structured correlations method [66], where one seeks to identify correlations between parameters and to calculate ranks (based on which parameters can be identified uniquely if they are not strongly correlated with other parameters). This approach utilises the total order sensitivity matrix to calculate these correlations. Therefore, the need for reliable and robust sensitivity matrices is vital to whichever method is implemented.

The work of Puy et al. [18] is conclusive in its findings of the Jansen and Janon estimators being the most reliable in finding the “truth” input parameter effects. Our work complements these conclusions in that we find the Jansen and Janon are the most reliable in the calculation of input parameter orthogonality, which appears to be motivated by input parameter convergence. Puy et al. [18] also reported that any choice made on the model have a non-negligible effect. While we agree with these conclusions for the most part, we are able to identify that similar to the model dimensionality, it appears choices we make, such as the sampling methodology and type of data used, have more impactful consequences. The lower variation associated with input parameter sets, when low discrepancy sequences are used, implies their effectiveness in returning robust and reliable input parameter sets. It appears that there is no clear advantage of using either continuous or discrete measurements when choosing how to calculate the total order indices.

## 5 Conclusion

Our study delved into the intricacies of varying sampling methodologies and variance-based total order estimators, aiming to establish best practices for practical identifiability studies. We conducted our investigation using two highly non-linear and stiff 0D models of the human cardiovascular system as our test cases: (i) a 1-chamber, 9-parameter model and (ii) a 2-chamber, 20-parameter model, both based on differential algebraic equations. Through a thorough empirical assessment of total order estimators and sampling methodologies, we gained valuable insights into their strengths and weaknesses, shedding light on the orthogonality of the input parameters within the models. This analysis complements prior work that focused on the estimators’ ability to uncover the “true” effects of a model, enriching our comprehension of their practical identification.

Our findings strongly advocate for the Jansen and Janon estimators as robust choices across different sampling methodologies, measurement data variations, and model dimensions. These two estimators emerge as preferred tools for calculating total order indices and, consequently, for identifying the optimal set of input parameters. Their efficient convergence and the consequential reduction in index uncertainty make them the optimal choice for this task. Furthermore, we recommend the use of low-discrepancy quasi-random Sobol and Lattice Rule sampling schemes as optimal sampling methodologies to complement Jansen and Janon estimators.

In essence, our work establishes a robust framework of good modelling practice for practical identifiability studies, considering both the influence of input parameters and their orthogonality. By incorporating these best practices into modeling studies, researchers can consistently and reliably identify the optimal input parameters for dynamical systems.

This approach not only enhances the quality and accuracy of parameter identification, but also paves the way for more informed decision-making in various scientific and practical domains.

## A Model Solutions and Parameters

In generic form, the equations relating to the passive compartmental state variables all take the form:

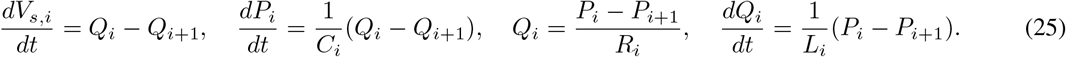

Above, the subscripts (*i* − 1), *i*, (*i* + 1) represent the proximal, present and distal system compartments, respectively. *V*_*s,i*_(mL) denotes the circulating (stressed) volume [67]. *C*_*i*_ (mL/mmHg), *R*_*i*_ (mmHgs/mL) and *L*_*i*_ (mmHg s^2^/mL) denote compartmental compliance, the Ohmic resistance and compartmental inertia between compartments *i* and (*i* + 1). See Figure 1 and Tables 14, 15.

**Table 14:** Input parameters for the 1 chamber model. Each input parameter’s unit is stated alongside a chosen initial value for the 9 parameter, 1-chamber model. *τ* is the cardiac cycle length and is fixed such that *τ* = 1*s*. The ventricular shift parameter *E*_shift_ = 0 s as no atrium is present in this model.

| Parameter $\theta$ (units) | Description | Initial Value |
| --- | --- | --- |
| $E_{max}$ $\left[ \frac{\text{mmHg}}{\text{ml}} \right]$ | Maximal ventricular contractility | 1.5 |
| $E_{min}$ $\left[ \frac{\text{mmHg}}{\text{ml}} \right]$ | Minimal ventricular contractility | 0.03 |
| $\tau_{es}$ (s) | End systolic time | $0.3\tau$ |
| $\tau_{ep}$ (s) | End pulse time | $0.45\tau$ |
| $Z_{ao}$ $\left[ \frac{\text{mmHg s}}{\text{ml}} \right]$ | Aortic valve resistance | 0.033 |
| $R_{mv}$ $\left[ \frac{\text{mmHg s}}{\text{ml}} \right]$ | Mitral valve resistance | 0.006 |
| $R_s$ $\left[ \frac{\text{mmHg s}}{\text{ml}} \right]$ | Systemic resistance | 1.11 |
| $C_{sa}$ $\left[ \frac{\text{ml}}{\text{mmHg}} \right]$ | Systemic compliance | 1.13 |
| $C_{sv}$ $\left[ \frac{\text{ml}}{\text{mmHg}} \right]$ | Venous compliance | 11.0 |

**Table 15:** Input parameters for the 2 chambers model. Each input parameter’s unit is stated alongside a chosen initial value for the 20 parameter, 2-chamber model. *τ* is the cardiac cycle length and is fixed such that *τ* = 1*s*. The ventricular shift parameter *E*_shift_ = 0.92 s as an atrium is present in this advanced 20 parameters model.

| Parameter $\theta$ (Units) | Description | Initial Value |
| --- | --- | --- |
| $E_{min_{lv}} \left[ \frac{\text{mmHg}}{\text{ml}} \right]$ | Minimal Ventricular Contractility | 0.1 |
| $E_{max_{lv}} \left[ \frac{\text{mmHg}}{\text{ml}} \right]$ | Maximal Ventricular Contractility | 2.5 |
| $\tau_{es_{lv}} \text{ (s)}$ | Ventricular Contraction | 0.3 |
| $\tau_{ep_{lv}} \text{ (s)}$ | Ventricular Relaxation | 0.45 |
| $E_{min_{la}} \left[ \frac{\text{mmHg}}{\text{ml}} \right]$ | Minimal Atrium Contractility | 0.15 |
| $E_{max_{la}} \left[ \frac{\text{mmHg}}{\text{ml}} \right]$ | Maximal Atrium Contractility | 0.25 |
| $\tau_{es_{la}} \text{ (s)}$ | Atrium Contraction | $0.045\tau$ |
| $\tau_{ep_{la}} \text{ (s)}$ | Atrium Relaxation | $0.09\tau$ |
| $Z_{ao} \left[ \frac{\text{mmHg s}}{\text{ml}} \right]$ | Aortic Valve Resistance | 0.033 |
| $R_{mv} \left[ \frac{\text{mmHg s}}{\text{ml}} \right]$ | Mitral Valve Resistance | 0.06 |
| $C_{sas} \left[ \frac{\text{ml}}{\text{mmHg}} \right]$ | Sinus Compliance | 0.08 |
| $R_{sas} \left[ \frac{\text{mmHg s}}{\text{ml}} \right]$ | Sinus Resistance | 0.06 |
| $L_{sas} \left[ \frac{\text{mmHg s}}{\text{ml}} \right]$ | Sinus Inertia | $6.2 \cdot 10^{-5}$ |
| $C_{sat} \left[ \frac{\text{ml}}{\text{mmHg}} \right]$ | Arterial Compliance | 1.6 |
| $R_{sat} \left[ \frac{\text{mmHg s}}{\text{ml}} \right]$ | Arterial Resistance | 0.05 |
| $L_{sat} \left[ \frac{\text{mmHg s}^2}{\text{ml}} \right]$ | Arterial Inertia | 0.0017 |
| $R_{sar} \left[ \frac{\text{mmHg s}}{\text{ml}} \right]$ | Arteriole Resistance | 0.5 |
| $R_{scp} \left[ \frac{\text{mmHg s}}{\text{ml}} \right]$ | Capillary Resistance | 0.52 |
| $R_{svn} \left[ \frac{\text{mmHg s}}{\text{ml}} \right]$ | Venous Resistance | 0.075 |
| $C_{svn} \left[ \frac{\text{ml}}{\text{mmHg}} \right]$ | Venous Compliance | 20.5 |

Figure 1 is a schematic representation for both the simple and advanced model. Note: (i) in Figure 1A, we use a C-R-C Windkessel [68] to represent the 2-chamber where as in Figure 1B, we use a C-R-L Windkessel to represent the aortic sinus and the systemic artery; (ii) no inertance appears in Figure 1A and there is no representation of the left atrium; (iii)all compartments in both models are passive, having fixed compliances; (iv) flow in and out of the active left atrium in Figure 1B is controlled by the systemic veins and the mitral valve. For both models, in Figures 1A and 1B, flow in and out of the active left ventricle is controlled by the mitral and aortic valves respectively. The valves are modelled as diodes, with Ohmic resistance under forward bias and infinite resistance under reverse bias:

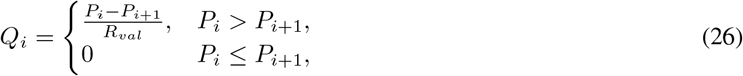

where *R*_*val*_ represents the resistance across the respective valve.

Let us consider the active model compartment. The dynamics of the left ventricle or left atrium are described by a time-varying compliance *C*(*t*), or reciprocal elastance, *E*(*t*) (mmHg/mL) which determines the change in pressure for a given change in the volume [67]:

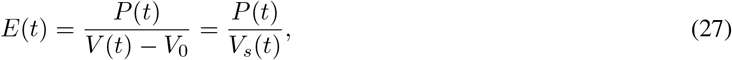

where *V*_0_ & *V*_*s*_(*t*) represent the unstressed and stressed volumes, respectively, in the left ventrical or left atrium. *E*(*t*) may be described in analytical form as follows: [49]

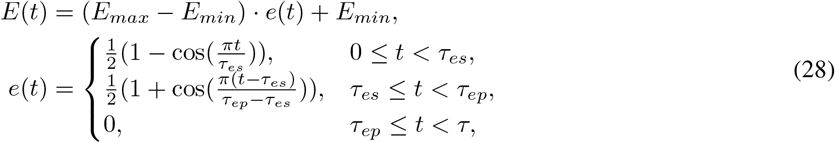

where *e*(*t*; *τ*_*es*_, *τ*_*ep*_) is the activation function for both the ventricle and the atrium and is parameterised by the end systolic and end pulse timing parameters *τ*_*es*_ and *τ*_*ep*_ respectively.

The elastance function is defined over one cardiac cycle, i.e., time 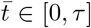 with *τ* (the length of the cardiac cycle) fixed in this work to *τ* = 1 s. The contractility, *E*_*max*_, and the compliance, *E*_*min*_, both control the elastance extrema of the left ventricular and the left atrium. There is a discontinuity in *E*(*t*) at *t* = *τ* when the next cycle starts.

Model A is implemented directly as a system of ODEs, Model B is implemented using an acausal modelling framework which will simplify the implementation of model variations. The acausal modelling library is published as a Julia package, CirculatorySystemModels.jl [53].

## Software & Data Availability

The analysis is performed with the programming language Julia [51]. All the analysis performed and example cases are freely available at https://github.com/H-Sax/Orthgonality-SA.

The acausal modelling library for Circulatory System Models is available at https://github.com/TS-CUBED/CirculatorySystemModels.jl and as a Julia package.

## Acknowledgments

Harry Saxton is supported by a Sheffield Hallam University Graduate Teaching Assistant PhD scholarship.

## Author Contributions

Conceptualisation: H.S,I.H,X.X, Formal Analysis: H.S,X.X, Methodology: H.S, X.X, T.S, I.H, Software: H.S, T.S, Supervision: I.H, X.X, R.C, T.S, Validation: X.X, I.H, Writing – Original Draft Preparation: H.S, I.H, X.X, Writing – Review & Editing: T.S, I.H, R.C, X.X

## Rights Retention Statement

For the purpose of open access, the authors have applied a Creative Commons Attribution (CC BY) licence to any Author Accepted Manuscript version arising from this submission.

## Footnotes

2 All other data is available at https://github.com/H-Sax/Orthgonality-SA

